# Environmental, host, and symbiont drivers of heat tolerance in a species complex of reef-building corals

**DOI:** 10.1101/2024.01.31.575130

**Authors:** MS Naugle, H Denis, VJL Mocellin, PW Laffy, I Popovic, LK Bay, EJ Howells

## Abstract

Reef-building coral populations are under unprecedented threat from climate warming. Yet, variation in coral heat tolerance exists whereby some colonies can cope with higher sea temperatures than others and thus may hold unique value for conservation and restoration. Here, we quantify variation in heat tolerance of an ecologically important tabular coral species complex across the Great Barrier Reef (GBR) while also measuring genomic variation in the coral host and symbiont partners. Coral bleaching and photochemical traits were measured in 569 colonies within the *Acropora hyacinthus* species complex from 17 reefs following exposure to standardized acute heat stress assays. We detected substantial variation in heat tolerance, where individual colony thermal thresholds differed by up to 7.3°C and 5.7°C among and within reefs, respectively. Sea surface temperature climatology was the strongest predictor of heat tolerance, where colonies from warmer northern and inshore reefs typically exhibited the highest thermal thresholds, while colonies from cooler southern reefs were able to tolerate greater temperature increases relative to their local summer temperatures. Heat tolerance was also positively associated with exposure to thermal stress in the weeks preceding measurements. Assignment of colonies to host genomic clusters revealed four putative species within the *A. hyacinthus* complex that did not vary in their responses to experimental heat stress. Symbiodiniaceae communities within colonies were comprised primarily of Cladocopium ITS2 variants that differed spatially but had minimal effect on heat tolerance. Between 36 - 80% of heat tolerance variation was explained by environmental, host, and symbiont genomic predictors, leaving 20 - 64% to be explained by additional underlying drivers such as functional genomic variation not measured here. These results may be used to inform conservation and restoration actions, including targeting heat tolerant individuals for selective breeding, and will provide a foundation for evaluating the genomic basis of heat tolerance.

## Introduction

Climate warming and unprecedented marine heatwaves have resulted in bleaching and mortality of reef-building corals (Glynn, 1996; Hughes et al., 2017; Leggat et al., 2019). However, not all corals will fair equally as sea temperatures continue to rise. Heat tolerance can vary widely among (Baird & Marshall, 2002; Evensen et al., 2022; Loya et al., 2001) and within coral species (Cornwell et al., 2021; Fuller et al., 2020; Humanes et al., 2022; Marhoefer et al., 2021). Intraspecific variation in coral heat tolerance may hold equal or even greater amounts of variation and ecological value compared to among-species variation, as has been shown in a wide variety of taxa and traits (Des Roches et al., 2018, 2021). Likewise, heat tolerance variation within closely related species complexes may be uniquely ecologically important, and especially valuable for adaptation if intraspecific variation is boosted through occasional hybridization between closely related taxa (Seehausen, 2004). Thus, understanding the heat tolerance repertoire of populations and closely related species is fundamental to understanding the future of coral reefs under climate change.

Intraspecific variation in the heat tolerance of corals is apparent among reef populations (Howells et al., 2013, 2016; Kenkel et al., 2013; Marzonie et al., 2023; Thomas et al., 2018), but can also occur within and among habitats across smaller spatial scales (Cornwell et al., 2021; Cunning et al., 2021; Humanes et al., 2022; Marhoefer et al., 2021). This phenotypic variation has been attributed in part to environmental factors such as thermal history, either by the adaptation through natural selection and/or through the acclimatization of individuals (Foo & Byrne, 2016; Palumbi et al., 2014; Thomas et al., 2018). Consequently, heat tolerance has been found to be positively associated with greater numbers of historical thermal anomalies (Marzonie et al., 2023; Quigley & van Oppen, 2022; Sully et al., 2019), or temperature variability (Carilli et al., 2012; Oliver & Palumbi, 2011). However, further studies are required to understand the distribution and drivers of intraspecific variation in heat tolerance at individual and ecosystem scales.

Quantifying intraspecific variation in key traits that affect fitness is important as it informs the potential for rapid evolution in response to environmental change. For example, intraspecific variation can affect community structure and dynamics, and thus is an important consideration for ecological or evolutionary models (Bolnick et al., 2011). Intraspecific variation may also be incorporated into coral projection models that calculate survival under future climate scenarios (Bay et al., 2017; Logan et al., 2014, 2021). Further, heat tolerant individuals are key targets for restoration and adaptation programs as they and their offspring may be more likely to persist under future warming scenarios than less tolerant conspecifics. Thus, a more comprehensive understanding of the drivers of variation in heat tolerance and the distribution and abundance of adaptive variation in reef metapopulations is crucial to inform evolutionary models, projections of future reef populations, selective breeding efforts, and conservation planning (Howells et al., 2022).

Coral heat tolerance may differ among closely related or cryptic species (Gómez-Corrales & Prada, 2020; Marzonie et al., 2023), including those within the *Acropora hyacinthus* complex, which is the focus of this study. *A. hyacinthus* represents a group of Indo-Pacific tabular corals (Ladner & Palumbi, 2012; Rose et al., 2021; Sheets et al., 2018; Suzuki et al., 2016) that hold exceptional value in ecosystem function but are threatened by a high vulnerability to climate warming (Ortiz et al., 2021). Although morphological differences exist among some taxa within this complex, it is often difficult or impossible to identify them without genetic data (Ladner & Palumbi, 2012). These cryptic lineages may vary in their growth rate, microhabitat distribution, and heat tolerance (Ladner & Palumbi, 2012; Rose et al., 2018, 2021; Suzuki et al., 2016).

Unresolved taxonomic diversity may lead to an overestimation of true intraspecific heat tolerance and may obscure investigations into drivers of heat tolerance. As species delineations within the *A. hyacinthus* complex is an active area of research (Ramírez-Portilla et al., 2022; Rose et al., 2021; Sheets et al., 2018), heat tolerance differences among putative species also need to be characterized. Currently, only two studies have assessed heat tolerance differences within species in the *A. hyacinthus* complex in American Samoa, and information is lacking for other regions including the GBR (Rose et al., 2018, 2021).

The community composition of algal endosymbionts (Symbiodiniaceae; LaJeunesse at al 2018) can also exert a strong influence on coral heat tolerance (reviewed in Quigley et al., 2018; van Oppen & Medina, 2020). Acroporid corals on the GBR have been shown to associate with primarily generalist *Cladocopium* species (Epstein et al., 2019; LaJeunesse et al., 2003; Matias et al., 2022; Oppen et al., 2005; Tonk et al., 2013), and associations with heat tolerant *Cladocopium* and *Durusdinium* species following bleaching have increased heat tolerance by over 1°C (Abrego et al., 2008; Berkelmans & van Oppen, 2006; Jones et al., 2008). Yet, heat tolerance conferred by Symbiodiniaceae associations may be species specific due to co-evolved symbiont-host partnerships and host-dependent effects (Abrego et al., 2008; Howells et al., 2012). Thus, to understand general patterns of heat tolerance conferred by Symbiodiniaceae associations, their effect on heat tolerance needs to be characterized in additional host species, including within the *A. hyacinthus* species complex.

Here, we quantify the heat tolerance traits of 583 colonies of *A. hyacinthus* from 17 reefs across 9.42° latitude of the GBR, in one of the largest-scale studies of intraspecific variation in coral heat tolerance to date. We use an acute heat stress assay to identify the spatial distribution of heat tolerant individuals and reef populations in a multi-metric approach. We then evaluate the relative influence of environmental factors, Symbiodiniaceae community, and host genetic identity on heat tolerance traits in this ecologically important species complex.

## Methods

### Field collections and acute heat stress assays

A total of 583 colonies of *Acropora hyacinthus* were sampled from 17 reefs encompassing a range of environmental conditions spanning over 1,200 km of the Great Barrier Reef (GBR; **Table S1**). Within reefs, colonies were primarily sampled from the reef crest or upper slope at a single site (i.e., within 5 km), apart from Hicks where a small number of colonies (n = 7) were sampled from an additional site. Since *A. hyacinthus* represents a species complex (Ladner & Palumbi, 2012; Ramírez-Portilla et al., 2022; Suzuki et al., 2016), the “neat” morphotype was targeted in this study to focus on intraspecific variation in heat tolerance. The “neat” morphotype is characterised by tightly packed and neatly arranged vertical branchlets with crowded labellate radial corallites forming a rosette-like arrangement, tightly reticulate basal branches, and a neater oval-shaped tabular structure (T. Bridge, pers comm). These characters distinguish it from similar species, where branch lengths and radial corallites are more irregular in shape and arrangement (T. Bridge, pers comm). However, a minority of colonies representing non-target morphotypes were also inadvertently sampled, particularly at sites where “neat” morphotypes were rarely encountered. The identity of colony morphotypes was confirmed by assignment to clusters based on genomic data (described below).

Colonies were sampled on SCUBA over the course of six field trips from January 2021 to March 2022. For each colony, 14 replicate fragments were collected; 12 fragments were placed in an acute heat stress assay, one was fixed in 100% ethanol for genomic analysis, and one provided a skeletal taxonomic reference. Colonies were sampled if they met size thresholds (> 20 cm) and showed no signs of disease or predation, though disease and predation were rarely encountered. Upon sampling, colony metadata was recorded including depth, GPS coordinates (as per Lukoschek et al., 2016), visual CoralWatch Health Chart Score (Siebeck et al., 2006), and photographs of each colony were taken for species identification. All colonies were sampled under permits from the Great Barrier Reef Marine Park Authority (G19/43148.1 and G21/45166.1) and with Free Prior and Informed Consent of Traditional Owners of the Sea Country where the study sites were located.

To quantify the heat tolerance of each colony within the *A. hyacinthus* complex, replicate fragments were exposed to a standardised acute heat stress assay (**Fig. 1**). This assay enables rapid phenotyping of numerous colonies and has been shown to provide rankings of heat tolerance that corresponded with rankings determined by traditional longer-term heat stress experiments in the Red Sea (Voolstra et al., 2020), and with recovery after heat stress (Walker et al., 2022). Acute heat stress assays were performed with treatments at each site’s Maximum Monthly Mean (MMM) and the MMM +3, +6, and +9°C (*n* = 3 fragments per colony per treatment). The MMM was calculated according to the protocol established by the National Oceanic and Atmospheric Administration (NOAA) and represents the upper limit of baseline historical temperatures (Liu et al., 2014). It is defined as the warmest sea surface temperature (SST) of the 12 monthly mean values for the years 1985-1990 and 1993 (Heron et al., 2015). The MMM is a widely used metric of coral climatology as it is used to calculate Degree Heating Weeks (DHW), an established metric to predict coral bleaching and compare heat tolerance (Heron et al., 2015). Significant coral bleaching is expected when DHW > 4°C-weeks and widespread bleaching and mortality is expected when DHW > 8°C-weeks (Eakin et al., 2010; Liu et al., 2003).

**Figure 1.**
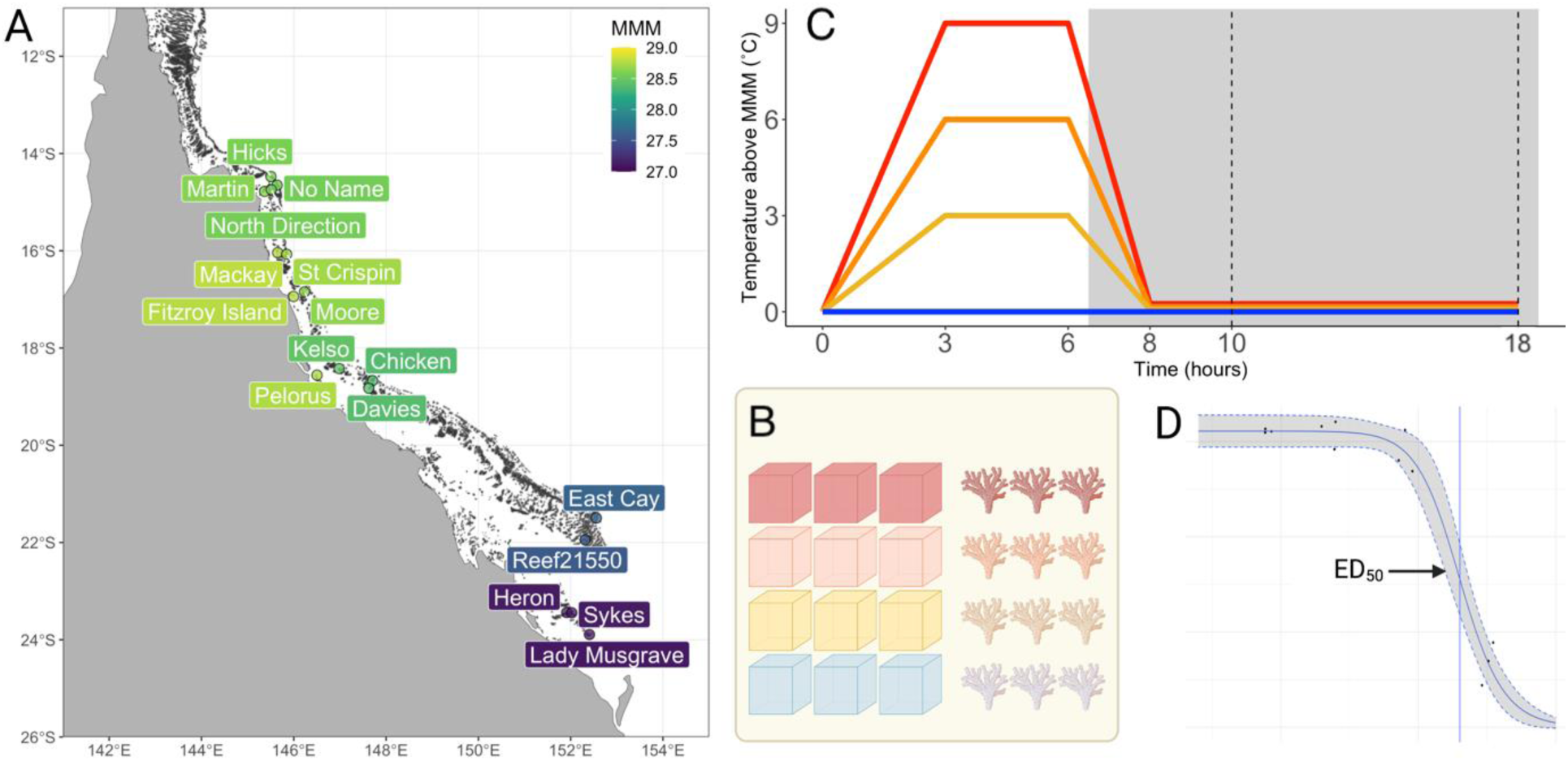
**A.)** Experimental design and heat tolerance trait data collection for *Acropora hyacinthus* colonies across the Great Barrier Reef. **B.)** Colonies were sampled from 17 sites on the GBR to be fragmented to allow 3 replicate fragments in each of 4 temperature treatments in experimental aquaria. **C.)** Aquaria were controlled to follow ramp-hold temperature profiles. Photochemical efficiency and chlorophyll content were measured after the ramp down from temperature (dashed lines, C). **D.)** Trait values at temperature treatments were used to calculate retained performance and fit log-logistic curves to generate ED_50_ values.

Heat stress assays were undertaken with a portable experimental system on board a research vessel at sea developed by the National Sea Simulator at the Australian Institute of Marine Science. The system had four independent temperature treatments with three tanks per treatment. Tanks were individually housed with exterior insulating jackets of seawater kept at treatment temperatures by independent heating elements (Omega 2 kW titanium) in separate sumps for each temperature treatment. Separate seawater lines passed through titanium coils (Waterco titanium heat exchanger), which proceeded through temperature-controlled sumps to reach their programmed temperature, and were directed into tanks at a flow rate of approximately 0.8L/min. Tanks contained temperature probes (TC Direct PT-100) and PAR sensors (Skye Quantum PAR – SKL26250) to maintain target temperatures and light levels.

Independent temperature loggers (HOBO) were also placed in select tanks to confirm appropriate temperature ramps after each run. Light levels were maintained at approximately 300 PAR (mmol photons/m^2^s^2^) and were gradually turned on and off at sunrise and sunset to mimic natural light conditions.

Fragments were mounted onto experimental racks immediately upon collection and held in flow-through aquaria overnight before being placed in their experimental tanks. Assay profiles involved a ramp up to treatment temperatures over 3 hours, a temperature hold for 3 hours, a ramp down to MMM over 2 hours, and an overnight hold at MMM. Profiles typically began at 11:00, with exceptions for Reef 21-550 (run 1 at 11:30 and run 2 at 12:00), Davies (12:00), East Cay (12:00), St. Crispin (run 1 at 12:30), and Martin (12:30). Downstream phenotypic measurements were delayed correspondingly to maintain consistent time spent in aquaria between the end of heat stress and trait measurements. One experimental assay has been removed from this dataset (run 2 at Reef 21-550) due to poor temperature control of aquaria during the experiment, reducing the final sample size to 569 colonies.

Additionally, heat tolerance was evaluated in 60 colonies within the *A. hyacinthus* complex at North Direction during a natural bleaching event in March 2021. Tolerance was measured using a CoralWatch Colour Health Chart Score to quantify bleaching (Siebeck et al., 2006) and a sample was preserved in 100% ethanol for genomic analyses. Since these colonies were found to encompass multiple putative species, they were used only for comparisons of natural bleaching responses among closely related putative species (described below).

### Measuring heat tolerance traits

Heat tolerance traits included photochemical efficiency (F_v_/F_m_) and chlorophyll content (NDVI), which were expressed as absolute thermal thresholds (i.e., the effective temperature inducing a 50% decline in trait performance; ED_50_), as well as their retained performance, (i.e., the proportion of retained trait performance under heat stress at +9°C above the local MMM). The photochemical performance of the symbionts within fragments was measured with an imaging PAM (Walz) two hours after the ramp-down to MMM was complete (dashed line, **Fig. 1**). Imaging PAM settings were as follows: measuring light intensity = 3 (in 2021) or 2 (in 2022, following replacement of LEDs), saturating pulse intensity = 7, gain = 1, damping = 1. From each fragment, three areas of interest were haphazardly selected, and the maximum quantum yield of photosystem II (F_v_/F_m_) was measured. The three F_v_/F_m_ values were averaged to attain a mean measurement per fragment. Inadvertently, imaging PAM chlorophyll fluorescence images were missing for one or two of the three replicate fragments for 6, 5, 1 and 47 of the 569 colonies at MMM, +3°C, +6°C, and +9°C respectively.

Chlorophyll content was measured using a hyperspectral camera (Resonon Pica XC2) ten hours after the ramp-down to MMM was complete (dashed line, **Fig. 1**). Three areas of interest were haphazardly selected on each fragment and a custom MATLAB script was used to extract a median Normalized Difference Vegetation Index (NDVI) per fragment. NDVI is an established metric to monitor land vegetation using satellites (Rouse et al 1973) but has also been used to measure chlorophyll a reflectance of soft corals in experimental aquaria (Leal et al., 2015; Rocha et al., 2015). Reflectance was normalized using a grey standard and NDVI was computed as (R_720_ – R_670_) / (R_720_ + R_670_). This approach was validated in our sample set by comparing hyperspectral-image generated NDVI values to total chlorophyll content in a subset of tissue samples from two reefs (Chicken and Davies). Total chlorophyll was extracted from homogenized powder and measured on a microplate spectrophotometer (Bio-tek Powerwave).

Chlorophyll was quantified and standardized to ash-free dry weight (Ritchie, 2006). NDVI values were highly correlated to total chlorophyll when compared among fragments (R = 0.74, p = <0.001, n = 293; **Fig. S1A**) and averaged across replicate fragments of each colony (R = 0.86, p <0.001, n = 98; **Fig. S1B**). This demonstrates that the NDVI trait captures the decline in chlorophyll that occurs with color paling during the bleaching stress response. NDVI metrics were not computed for colonies in the first run at Hicks (n = 20 colonies) or the first run at Reef 21-550 (n = 15 colonies), as hyperspectral images were not taken.

From each of these two phenotypic traits (F_v_/F_m_ and NDVI), two heat tolerance metrics were computed: an absolute metric which represents an absolute thermal threshold independent of local thermal history and a retained heat tolerance metric that represents how traits decline relative to their local site MMM. Absolute heat tolerance was measured as the effective dose (ED) temperature (°C) at which trait performance declines by 50% (i.e., ED_50_). ED_50_ values were estimated by fitting three-parameter log-logistic dose response curves in the `drc` package in R with the following constraints: slope = [10,120]; ED_50_ = [28,40], F_v_/F_m_ = [0,0.8], NDVI = [0,1]. ED_50_ values with confidence intervals greater than 5°C were excluded, and 83.7% and 91.8% of colony values were retained for downstream analyses for F_v_/F_m_ and NDVI, respectively. Retained heat tolerance was quantified as the proportion of trait performance retained at the highest heat stress treatment (MMM+9°C). Metric values were corrected for any deviation between target treatment and actual tank temperatures, such that retained heat tolerance = (TraitMMM+9°C / TraitMMM) * ((TempMMM+9°C – TempMMM) / 9).

Variation within and among sample sites was investigated using linear mixed models in the package `lme4` in R (Bates et al., 2015). In the first iteration, site was set as a random factor, with residual variance attributed to within-site variation. In a second iteration, site and genomic cluster (described below) were each set as random factors to assess residual variation within both sites and genomic clusters.

### Genomic assignment to divergent species clusters

Genomic DNA was extracted using the QIAGEN Blood and Tissue kit and purified with SPRI magnetic beads following manufacturers specifications. Whole genome libraries were prepared using the Lotus DNA Library Prep Kit for NGS with 10 ng of input DNA and enzymatic fragmentation to achieve average insert sizes of 350 bp. Final amplification consisted of 8 PCR cycles. Libraries were quantified using a Quant-iT dsDNA assay kit and up to 192 libraries were multiplexed in equimolar ratios for sequencing to achieve 10x coverage. Sequencing was performed by Azenta Life Sciences on the NovaSeq S4 300 cycle using 150 bp paired end reads. Raw sequence data are available under the NCBI Sequence Read Archive: PRJNA982441.

`FASTQC` (http://www.bioinformatics.bbsrc.ac.uk/projects/fastqc) and `MultiQC` (Ewels et al., 2016) were used to examine read quality and adapter contamination for each sample. Quality trimming of raw reads was performed in Trimmomatic version 0.39 (Bolger et al., 2014) using a 4 bp sliding window, a minimum phred score quality of 20, and a minimum read length of 50 bp. Adapter sequences were removed using the Illuminaclip option in ‘palindrome mode’. Trimmed reads were mapped to the *A. hyacinthus* genome (López-Nandam et al., 2023) using the Burrow-Wheeler Aligner (Li & Durbin, 2009) version 0.7.17 and the MEM algorithm with default settings. The resulting alignment SAM files were converted to indexed and sorted BAM files using `Samtools` v1.10 (Danecek et al., 2021). PCR duplicates were removed using `picard` (http://broadinstitute.github.io/picard/). Individuals with less than 80% of mapped reads to the reference genome were excluded from downstream analyses. We also removed technical replicates, retaining a total of 565 samples for which we had viable genomic and phenotype data and 60 colonies collected during a natural bleaching event.

Genomic data was visualized using `ANGSD` version 0.934 (Korneliussen et al., 2014) to identify genetically distinct host clusters. We identified polymorphic sites and estimated genotype likelihoods to account for statistical uncertainty associated with sequencing errors or missing genotypes. Polymorphic sites were filtered as: mapping quality > 30, base quality > 30, coverage ≥ 3 reads in at least 95% of individuals, and sites mapping to 14 assembled chromosomes. We called major and minor alleles directly from the genotype likelihoods assuming biallelic sites and considered only polymorphic sites with a likelihood ratio test p-value <0.000001. To evaluate population genetic structure, we extracted an individual covariance matrix with `PCAngsd` (Meisner & Albrechtsen, 2018) applying a 0.05 minor allele frequency (MAF) threshold. We computed eigenvectors and eigenvalues in R and performed a Principal Components Analysis (PCA). We also performed Bayesian hierarchical clustering admixture analyses in `PCAngsd` using the ‘admix’ option to estimate individual ancestry proportions assuming 2-5 ancestral populations (K genetic clusters) and applying a MAF threshold of 0.05 (Meisner & Albrechtsen, 2018).

### Quantifying Symbiodiniaceae communities

Symbiodiniaceae communities were quantified using the ITS2 rDNA marker (Hume et al., 2019) using established protocols for PCR amplification, library preparation, and ITS2 sequencing (see Supplementary Methods and Results).

The SymPortal framework (https://symportal.org/; Hume et al., 2019) was used to assign reads within each sample to Symbiodiniaceae genera, subgeneric ITS2-type profiles, and unique sequence variants. The Symbiodiniaceae ITS2 rDNA region can resolve genera and some species-level variation, however closely related species and sub-species can be challenging to delineate due to high intragenomic sequence diversity (Davies et al., 2022). The SymPortal framework addresses this issue by assigning ITS2-type profiles to sets of co-occurring sequence variants which are more likely to belong to the same genotype when they repeatedly co-occur in biological samples (Hume et al., 2019). Therefore, ITS2-type profiles represent putative Symbiodiniaceae taxa while sequence variants represent intragenomic variation that may or may not confer phenotypic or species differences (Davies et al., 2023). ITS2 reads were normalized by median sequencing depth and sequences with fewer than 10 reads were removed, resulting in a total of 52 unique sequence variants across the sample set. ITS2-type profiles were also removed if they contained fewer than 10 reads, resulting in 12 ITS2-type profiles. To reduce the prevalence of metabarcoding artifacts, samples with low overall read counts were discarded from the analysis. Minimum read count thresholds were set as the number of reads found in negative control samples: 7,500 reads for ITS2 sequence variants and 6,100 reads for ITS2-type profiles. This reduced the sample size to 424 colonies with assigned ITS2-type profiles and 421 colonies with ITS2 sequence variant data.

To characterize the Symbiodiniaceae community composition in each *A. hyacinthus* colony, the dominant ITS2-type profile was examined since over 97% of samples were assigned a single profile. However, as ITS2-type profile assignments in *A. hyacinthus* on the GBR have not yet been validated with other genetic makers, we also examine the relative abundance of all ITS2 amplicon sequence variants (ASVs) within our samples. These were summarized using Principal Components (PC) values of normalized read counts of all ITS2 sequences. Both ITS2-type profiles and ITS2 sequence variants (expressed as PCs) were examined for their relationships to shelf position, latitude, host genomic cluster, and their interactions using PERMANOVA on Bray-Curtis dissimilarities with 999 permutations. This was performed with the adonis2 function within the `vegan` package in R (Oksanen et al., 2022).

### Environmental variables

A total of 22 environmental variables were computed to quantify site- and colony-level variation in thermal history, accumulated heat stress, nutrient availability, and light penetration (**Table S2**). This included data obtained from NOAA CoralWatch (Liu et al., 2014), eReefs (Steven et al., 2019), and NASA MODIS (Savtchenko et al., 2004), as well as in situ measurements (i.e., colony depth and pigmentation) (**Table S2**). Variables computed from eReefs are based on hydrodynamic or biogeochemical models that estimate variables at different vertical stratifications. Thus, eReefs-derived variables were computed using values from models at the closest vertical stratification to the true colony depth (from -0.5, -2.35, -5.35, -9, or -13 m for the GBR1_hydro hydrodynamic model and -0.5, -1.5, -3, -5.5, -8.8, -12.75 m for the GBR4_BGC biogeochemical model). Additionally, some temperature variables were calculated using eReefs while others were calculated using CoralWatch, since these datasets vary in their spatial resolution and temporal range. Variables of climatology and thermal anomalies were computed using CoralWatch which ranges from 1985 and has a 5 km resolution, while variables of warming rate and temperature variability were computed using eReefs which ranges from 2014 and has a 1 km resolution (**Table S2**).

### Gradient Boosting and Prediction Models

Predictors encapsulating variation in environment, endosymbiont community, and host genomic cluster were examined for their influence on heat tolerance traits using boosted regression trees (BRT; **Table S2**). BRTs are a machine learning algorithm that evaluates the relationship between predictor variables and outcomes by building decision trees, that are boosted such that after each tree is built, a subsequent tree is trained using the residuals not explained by the previous tree (Elith et al., 2008). BRT algorithms work best when they learn slowly, building many trees and then averaging the patterns across all the trees to determine the influence and effect of predictor variables (Elith et al., 2008). BRTs can accommodate continuous and categorical variables and are robust to non-linear interactions and missing data (Elith et al., 2008). Collinear predictor variables do not affect predictive accuracy computation but may compete within the BRT model such that collinear variables may be elevated or repressed in variable importance rankings. Thus, the favored collinear variable in a BRT is more likely to be related to the response than its collinear predictors which were repressed. BRTs were implemented as gradient boosting machine models in the R package `gbm`. Model parameters were chosen as follows: [Learning rate = 0.001, Interaction depth = 5, Bag fraction = 0.5, CV folds = 10]. Diagnostic plots of holdout deviance by tree iteration were used to assess overfitting. To assess deviance explained by each model, an identical model was fit using the gbm.step function in the `dismo` package and was calculated as: (total deviance – cross validated residual deviance) / total deviance (Hijmans et al., 2013; Leathwick et al., 2006). Linear models and stepwise multiple regression were also used to investigate effects of predictors on heat tolerance traits and are explained in further detail in the Supplementary Methods and Results. All analyses were performed in R version 1.3.1093 and can be found on Github: https://github.com/melissanaugle/Ahya_heat_tol_variation

## Results

### Variation in heat tolerance across the Great Barrier Reef

Responses of colonies within the *A. hyacinthus* complex to standardised acute heat stress assays varied considerably across the GBR (**Fig. 2**). Overall mean ED_50_ temperatures were 36.4°C and 35.6°C for F_v_/F_m_ and NDVI respectively, and these differed among colonies by 3.5°C (F_v_/F_m_: 34.6-38.1) and 7.3°C (NDVI: 32.4-39.7; **Table 1**). For both traits, colonies with the highest ED_50_ temperatures predominantly occupied northern reefs, however a few individuals ranked within the top 20% were also found at central and southern reefs (**Fig. 2A-B**). Mean retained performance values were 0.39 and 0.26 for F_v_/F_m_ and NDVI respectively, ranging by 0.93 (F_v_/F_m_: 0.00-0.93) and 0.91 (NDVI: 0.03-0.95). Colonies with the highest retained performance generally occupied southern reefs with some individuals within the top 20% also occupying central and northern reefs, a pattern that was stronger for retained F_v_/F_m_ (**Fig. 2B**) than retained NDVI (**Fig. 2C**).

**Figure 2.**
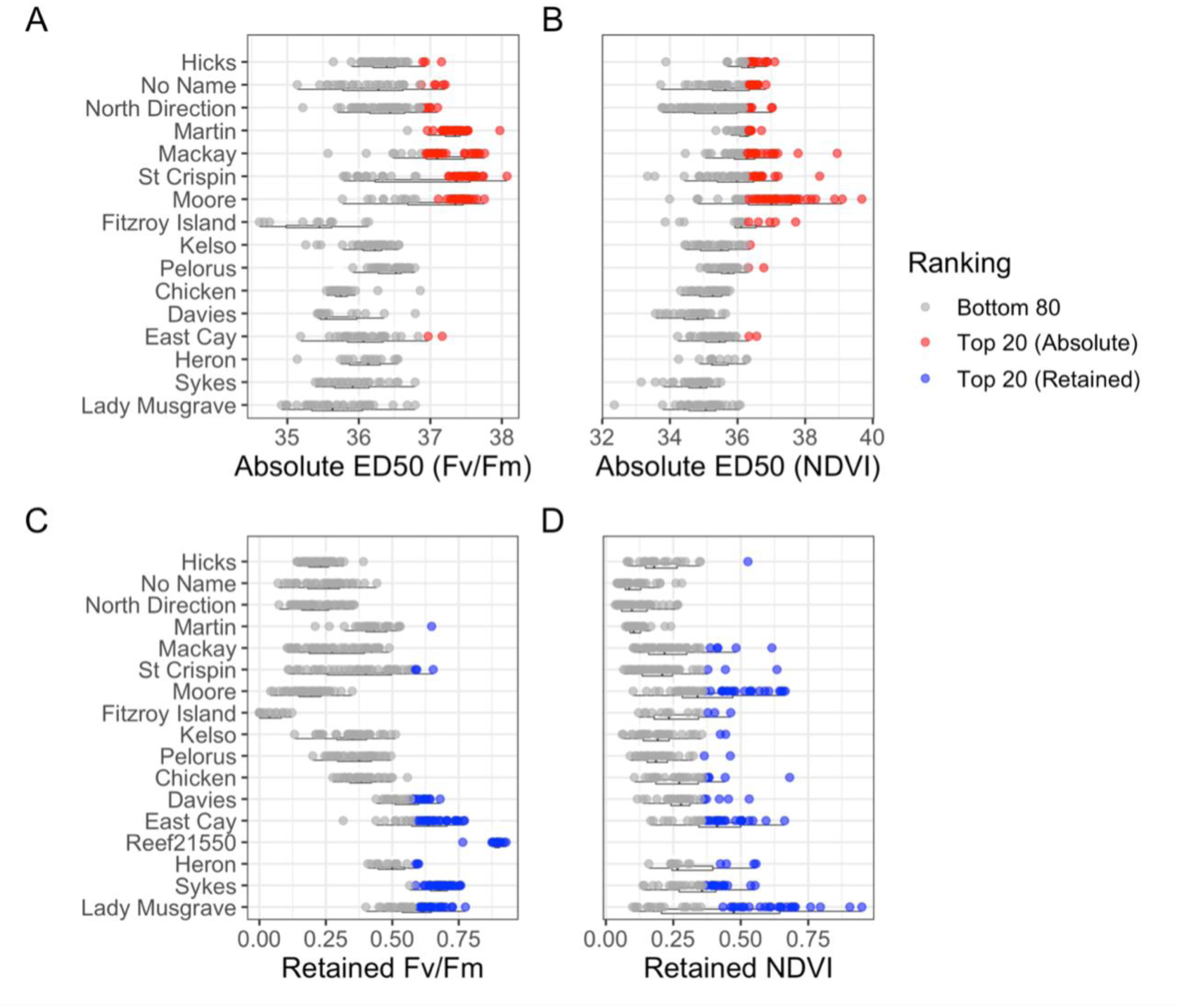
Variation in heat tolerance trait metrics in *Acropora hyacinthus* colonies across sites on the Great Barrier Reef organized from north to south. For absolute metrics (i.e., ED_50_ thresholds), of the maximum quantum yield of photosystem II (F_v_/F_m_, **A**) and the normalised difference vegetation index (NDVI, **B**), most heat tolerant individuals (i.e., within the top 20%) were from the northern GBR. In contrast, for retained performance metrics of F_v_/F_m_ (**C**) and NDVI (**D**), most heat tolerant individuals were from the southern GBR. The distribution of top performing colonies also varied by metric, whereby top performers according to F_v_/F_m_ were found in a smaller number of sites and top performers according to NDVI can be found across most sites.

**Table 1.**
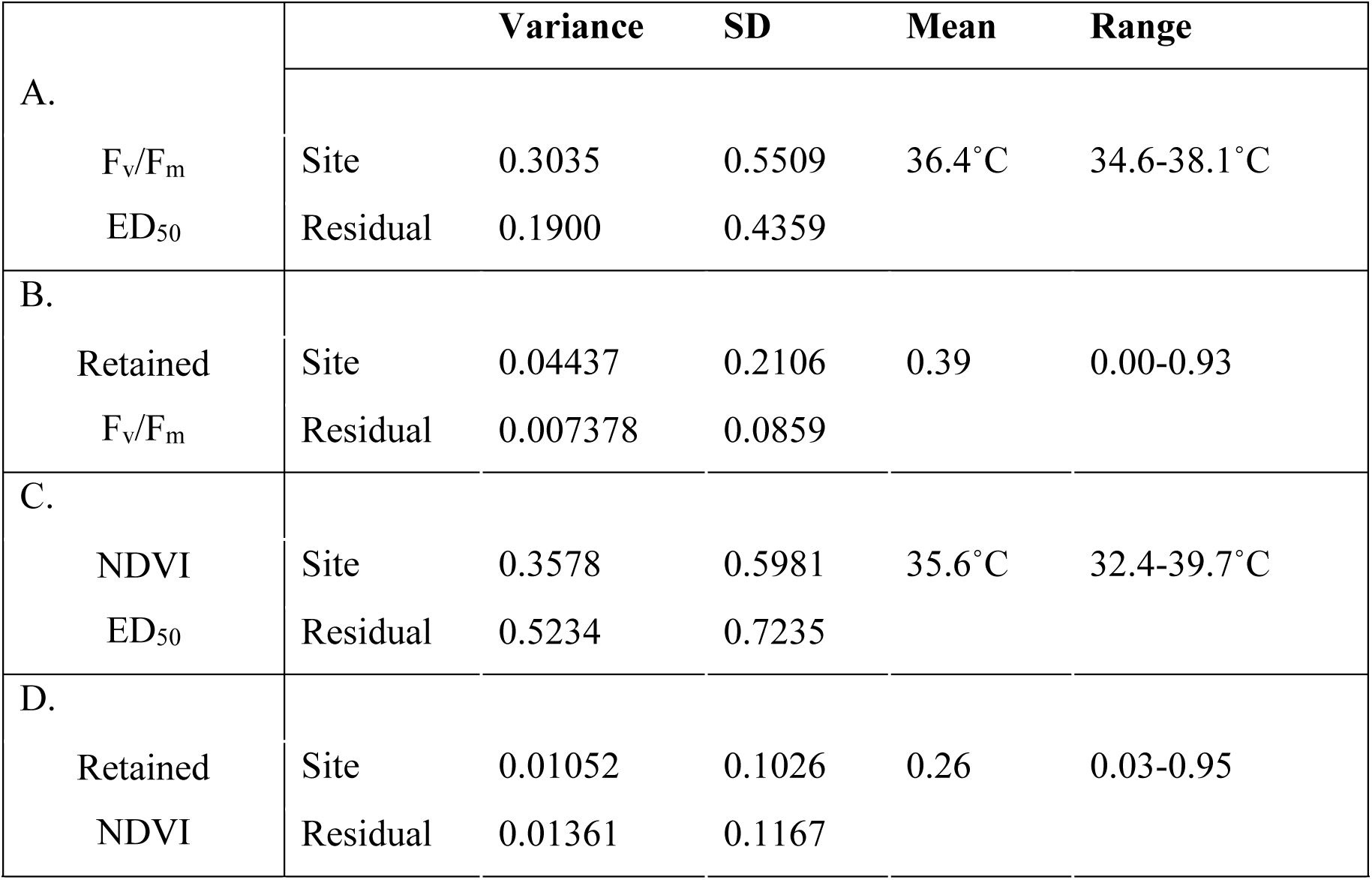
Linear mixed effects models (LMMs) showing the among-site variation and residual (within-site) variation, as well as mean and range in values, of heat tolerance trait metrics in *Acropora hyacinthus* colonies across sites on the Great Barrier Reef. Greater among site variance was exhibited by the maximum quantum yield of photosystem II (F_v_/F_m_) absolute metric (ED_50_ threshold; **A**) and retained performance metric (**B**), while greater within-site variance was exhibited by the normalised difference vegetation index (NDVI) absolute metric (ED_50_ threshold; **C**) and retained performance metric (**D**).

Within-site rankings were largely consistent between absolute and retained performance metrics of heat tolerance for both photochemical capacity (F_v_/F_m_; *R*=0.71) and chlorophyll content (NDVI; *R*=0.65; **Fig. S2**). The relationship of tolerance rankings between F_v_/F_m_ and NDVI traits was weaker (*R* = 0.36-0.47; **Fig. S3**), potentially as these traits capture different biological aspects of the heat stress response with F_v_/F_m_ capturing a breakdown in the photochemical pathways within Symbiodiniaceae and NDVI capturing the loss of Symbiodiniaceae during bleaching. Consequently, trade-offs existed between heat tolerance responses across the GBR, where few colonies performed highly for both ED_50_ and retained performance metrics and tended to be restricted to northern-central and southern reefs, respectively (**Fig. 2**).

For ED_50_ and retained metrics of F_v_/F_m_, variance was 1.6 – 6 times higher among sites than within sites (**Table 1**). However, for both metrics of NDVI, variance was 1.3 – 1.5 times higher within sites than among sites, and this trend was most pronounced for ED_50_ thresholds (**Table 1**). Variance within and among sites for all heat tolerance metrics deviated by < 0.01 from variances stated above when also accounting for genomic cluster as a random factor (**Table S3**).

### Host genomic cluster assignments

For 625 samples, including those placed in the acute heat assay (n=565) and those assessed for their response natural bleaching (n=60), the number of quality filtered sequence reads ranged between 15,603,604 to 63,841,053 reads per sample, with a mean of 26,493,269 reads. Sequenced samples had an average of 92.5% primary reads mapped to the reference genome. The mean coverage among individuals was 11.4x genome-wide and 7.02x considering only scaffolds corresponding to 14 assembled chromosomes.

Principal components analysis of 1,238,851 loci in PCAngsd indicated that sampled coral colonies belonged to four distinct genetic clusters with 4.64% and 1.14% of genetic variation explained on the first and second PCs (**Fig. 3A**). The assignment of four major genetic clusters was also supported in Bayesian clustering analysis (**Fig. 3C**). Most colonies (472 of 625) were assigned to Group 1 (**Grey, Fig. 3A**) which represents our target morphotype, *A. hyacinthus* “*neat*”. Groups 2 – 4 contained 27, 36, and 90 colonies, respectively. Group 1 differentiated from Groups 2 - 4 across PC1 and its divergence was also supported in Bayesian admixture analyses for all values of K (**Fig. 3C**). Groups 2 to 4 likely represent other putative species (or ecotypes) within the *A. hyacinthus* species complex. Colony morphology and branchlet structure varied among genomic clusters (**Fig. 3B**), yet colonies of the same cluster often presented variable features in different sites and may be very difficult to distinguish based on morphology alone (i.e., cryptic).

**Figure 3.**
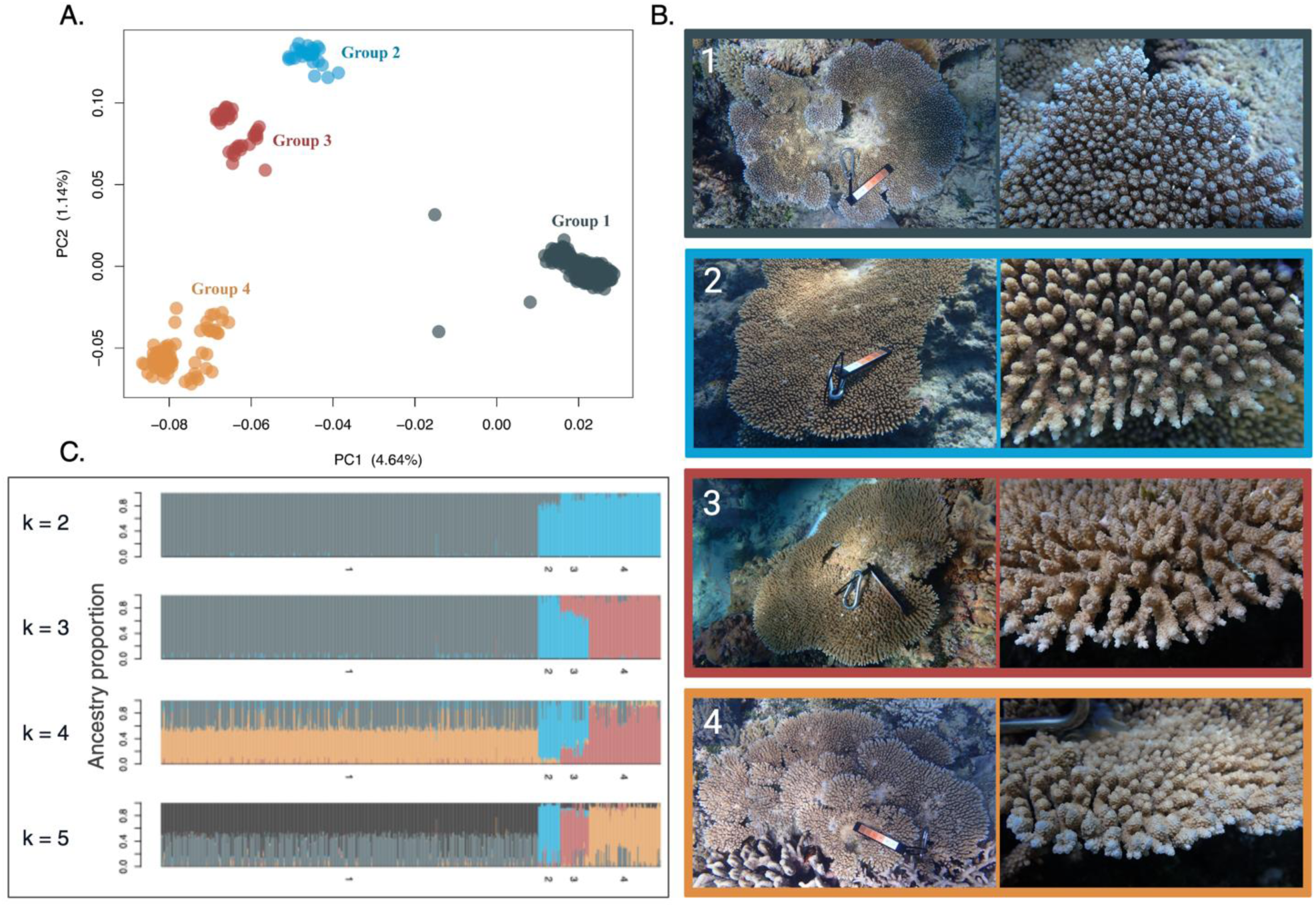
**A.)** Principal Components Analysis of 1,308,950 SNPs showing four distinct clusters representing four distinct putative species within the *Acropora hyacinthus* complex, sampled across the Great Barrier Reef. Group 1 represents the target species, *A. hyacinthus* “*neat*” which comprised the majority of colonies sampled (472 of 625 total colonies), with groups 2 - 4 containing 27, 36 and 90 colonies, respectively. **B.)** Colony morphology and branching structure of genomic clusters 1 - 4 within the *A. hyacinthus* species complex on the Great Barrier Reef. Colonies pictured to represent clusters 1-4 were all sampled on Sykes reef. Clusters were assigned based on differences in genome-wide SNPs. Host genomic cluster was assigned based on genome-wide SNPs with group 1 representing *A. hyacinthus* “neat” and groups 2 - 4 representing other putative species in the *A. hyacinthus* complex. **C.)** Admixture plot assuming 2 - 5 ancestral populations (k) within the *A. hyacinthus* species complex sampled across the Great Barrier Reef, supporting the grouping of these colonies into four putative species clusters.

Host genomic clusters (**Fig. 3A**) varied by site, with Group 1 (*A. hyacinthus* “*neat*”) sampled primarily in the central and northern GBR and Groups 2 - 4 sampled primarily in the southern GBR and at Fitzroy Island (inshore northern GBR; **Fig. 4A**). While differences may exist in the distribution and abundance of these genomic clusters (or putative species), the relationship between latitude and cluster prevalence seen here could also reflect collection bias and identification error at different sites. Thus, we restricted our analysis of heat tolerance variation among clusters to the 5 southern sites to reduce the confounding latitudinal effects on heat tolerance (Groups 1 - 4: n = 44, 14, 27, 51). Site-adjusted heat tolerance metrics exhibited higher variation within than among genomic clusters (**Fig. 4B-E**), although Group 2 displayed a non-significant trend for lower NDVI ED_50_ (**Fig. 4D**). This relationship was more pronounced during a natural bleaching event, where Group 2 and 3 exhibited lower CoralWatch Colour Scores than Group 4, indicating a lighter appearance and greater bleaching sensitivity of these putative species (**Fig. S4**).

**Figure 4.**
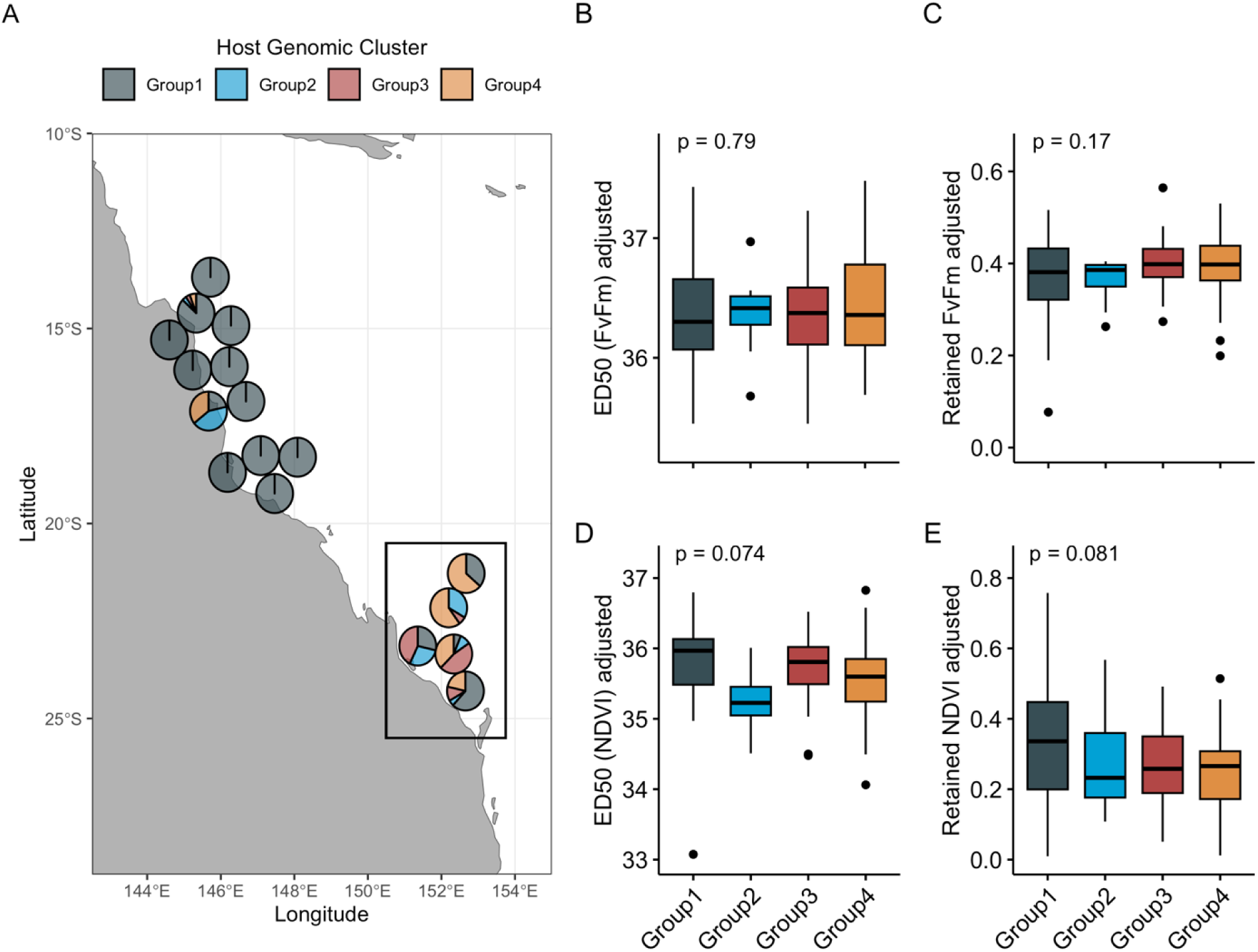
**A.)** Variation in *Acropora hyacinthus* host genomic cluster assignments across sites on Great Barrier Reef. **B - E.)** Heat tolerance traits from the southern reefs (sites shown in rectangle in A). Groups 1 - 4 with from southern reefs shown in rectangle contained 44, 14, 27, and 51 individuals, respectively. Host genomic clusters were assigned based on genome-wide SNPs with Group 1 representing *A. hyacinthus* “neat” and Groups 2 - 4 representing other putative species in the *A. hyacinthus* complex. Heat tolerance traits have been adjusted to remove variation due to site using residuals from site-specific means. P-values are shown for each trait using one-way ANOVAs.

### Symbiodiniaceae communities

Symbiodiniaceae ITS2 communities in *A. hyacinthus* colonies were overwhelmingly composed of *Cladocopium* variants (≥ 99% of sequencing reads in all colonies; **Fig. 5B**), with rare occurrences of *Symbiodinium* (4% of reads in 22% of colonies). The major *Cladocopium* variants in our sample set co-occurred within colonies across the GBR, indicating they are of intragenomic origin. Variants differed spatially, most remarkably across the inshore-offshore gradient (**Fig. 5A**). Two closely related variants, C50a and C50c were disproportionately detected in inshore sites (yellow, **Fig. 5B**), while C50b was detected primarily in southern sites (purple, **Fig. 5B**). The C3k variant was detected in every colony (n = 421), although northern mid and outer reef sites contained proportionally more C3k reads compared to southern and inshore sites. Consequently, 70% of colonies were assigned a single dominant ITS2-type profile (C3k/C50a/C50b/C50c-C3ba-C50q-C3-C50f) which occurred at all reefs but was hosted almost exclusively on inshore reefs (blue, **Fig. S5**). A further 16% were assigned a closely related ITS2-type profile (C3k-C50a-C3-C3ba-C50q-C50f-C3dq-C3a), which was most common on mid and offshore reefs (yellow, **Fig. S5**). The remaining 9 ITS2-type profiles were each hosted by less than 4% of colonies, with C3k/C3bo-C3ba-C50a-C50q-C50b occurring primarily on southern reefs (Purple, **Fig. S5**).

**Figure 5.**
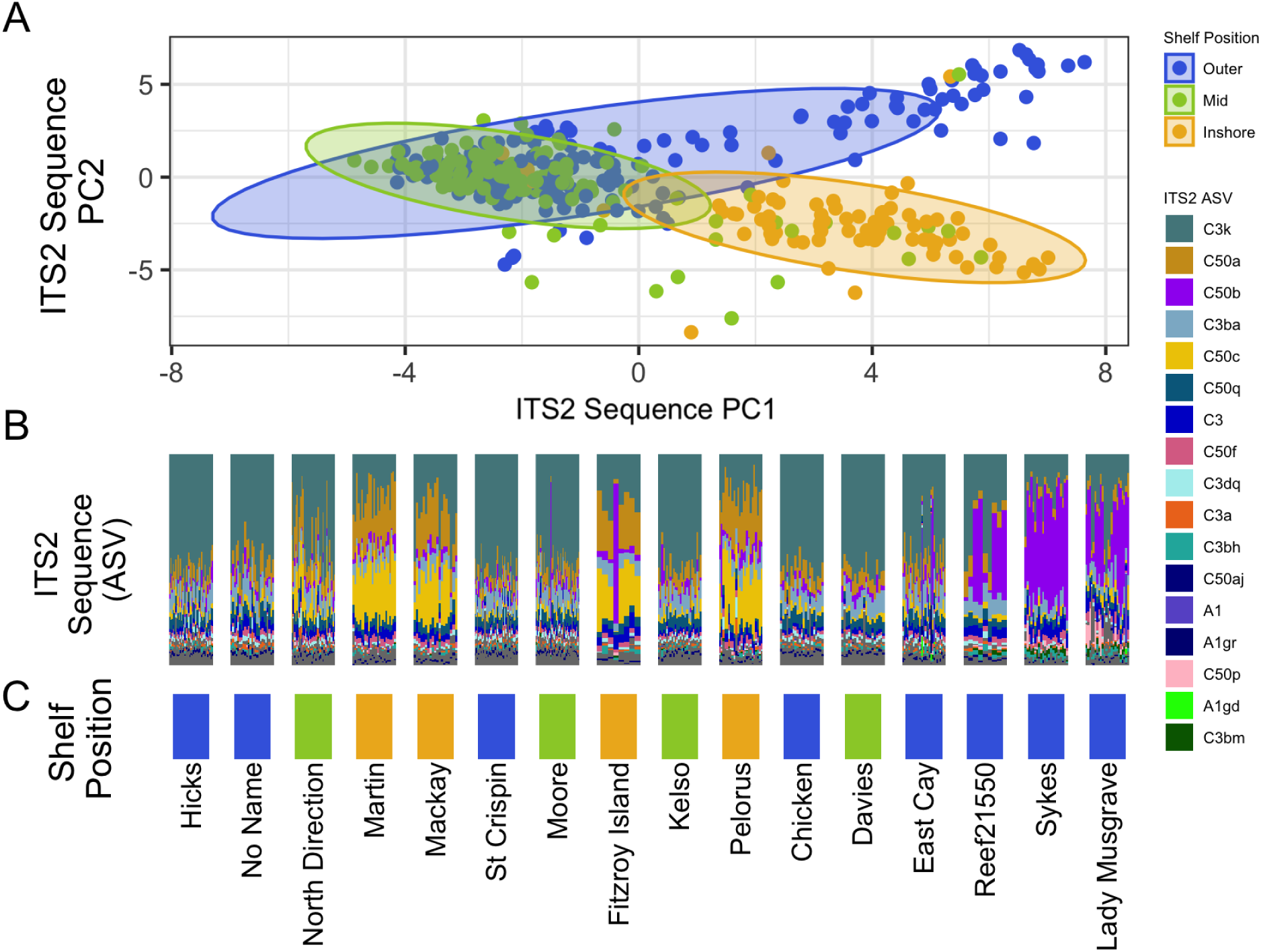
**A.)** Principal components analysis (PCA) of ITS2 sequences in 404 colonies of *Acropora hyacinthus* across the Great Barrier Reef showing ITS2 sequence differentiation between colonies in inshore habitats versus those in outer and mid-shelf habitats. **B.)** Variation in Symbiodiniaceae communities within *A. hyacinthus* colonies across the Great Barrier Reef. ITS2 sequences are shown for individuals (vertical columns) within sites (arranged by latitude). **C.)** Shelf position (bottom row) is shown to visualize ITS2 sequence differences at inshore sites (yellow, bottom row).

ITS2-type profiles and sequences were most strongly differentiated across shelf position and latitude; a trend that was 10 times more pronounced for ITS2 sequence variants (PERMANOVA, partial ω^2^ = 0.41 and 0.23, p = 0.001; **Table S4**) than type profiles (PERMANOVA, partial ω^2^ = 0.035 and 0.026, p = 0.001; **Table S5**). ITS2 sequences revealed similar trends between offshore and mid-shelf sites, with more distinct sequences found in inshore sites (**Fig. 5A**) While we did not detect variation in ITS2-type profiles by host genomic cluster, there were minor differences in sequence variants among host genomic clusters (PERMANOVA, partial ω^2^ = 0.090, p = 0.001), that interacted with the effects of shelf position and latitude (PERMANOVA, partial ω^2^ = 0.027 and 0.091, p = 0.001; **Table S4**).

Despite this spatial variation, we did not observe any relationship between the first PC value summarizing Symbiodiniaceae ITS2 sequence variants in *A. hyacinthus* and heat tolerance metrics across the GBR, apart from a very weak relationship with F_v_/F_m_ ED_50_ (*R* = -0.12, p = 0.026; **Fig. S6**). This was also the case at the site-level, apart from F_v_/F_m_ ED_50_ at North Direction (*R* = -0.39, p = 0.019, **Fig. S7**) and both F_v_/F_m_ metrics at Lady Musgrave (*R* = -0.45 and -0.51; p = 0.029 and 0.027; **Fig. S7**).

### Top predictors of heat tolerance traits

Boosted Regression Trees (BRTs), incorporating environmental, Symbiodiniaceae, and host genomic cluster predictors, explained 35.9% to 80.6% of the variation in heat tolerance of *A. hyacinthus* across the GBR (**Fig. 6**). Temperature variables were the most predictive of trait variation for all metrics. MMM related variables held strong relative predictive influence for both ED_50_ metrics whereby a MMM > ∼28.5°C signalled an increase in F_v_/F_m_ and NDVI ED50s (**Fig. 6A,C**). However, MMM showed a negative relationship with retained performance, indicating that colonies on cooler reefs are living further from their upper thermal limits (**Fig. S8**). F_v_/F_m_ metrics tended to have stronger relationships to MMM than NDVI metrics, likely because F_v_/F_m_ primarily varied across sites while NDVI varied more strongly within sites.

**Figure 6.**
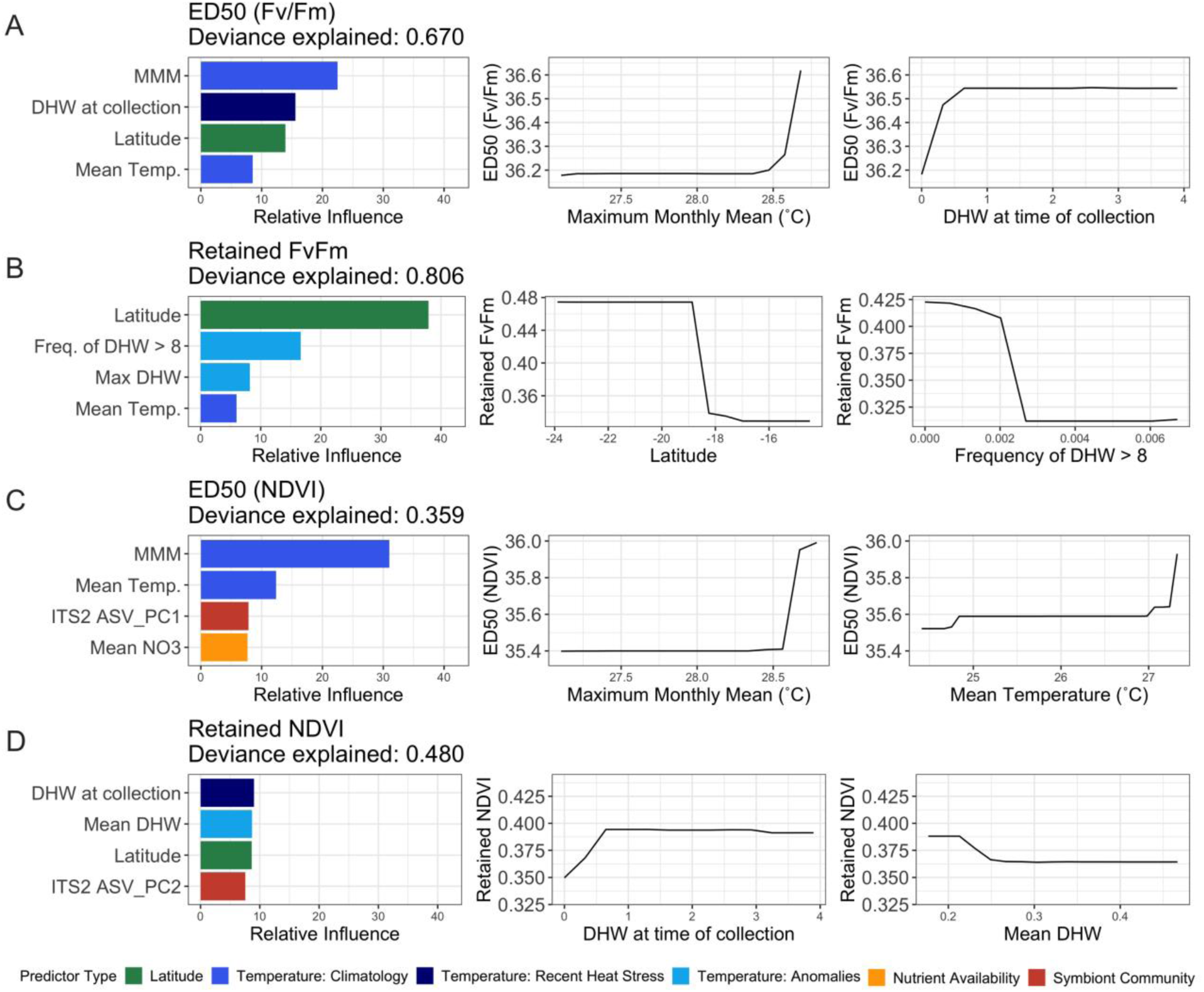
Boosted regression tree model outputs showing the relative influence of environmental, Symbiodiniaceae, and host genetic identity predictors on heat tolerance traits of *Acropora hyacinthus* on the Great Barrier Reef including **A.)** F_v_/F_m_ ED50, **B.)** retained F_v_/F_m_, **C.)** NDVI ED_50_ and **D.)** retained NDVI. Plots show the top four predictors for each heat tolerance metric (left column) and the marginal effect of features of interest on heat tolerance (right columns).

Degree Heating Weeks (DHW) at the time of collection was an important predictor for 2 of 4 metrics, F_v_/F_m_ ED_50_ and retained NDVI, indicating that exposure to recent heat stress (DHW > 0.5) improved performance in our heat stress assays (**Fig. 6A, D**). Thermal anomalies were also an important predictor for retained metrics, where performance was highest at southern reefs that have experienced fewer thermal anomalies (**Fig. 6B, D**). Symbiodiniaceae variables appeared in the top 4 predictors for NDVI metrics (**Fig. 6C, D**), and host genomic cluster was not a top predictor in any of the BRT models.

BRT models of retained performance metrics explained 12.1-13.2% more variation than absolute (ED_50_) metrics, likely because retained traits varied more among sites and ED_50_s varied more within sites, and most environmental predictors were stratified at the site-level.

Additionally, predictors explained 31.1-32.6% more variation in F_v_/F_m_ than NDVI, likely due to the high within-site variability of NDVI metrics. These results were supported by linear models which showed that the amount of variance explained by predictors and traits was largely consistent with BRTs (**Fig. S9**; **Table S6**).

## Discussion

Our study provides one of the largest-scale investigations of intraspecific variation in the heat tolerance of reef-building corals. Using acute heat stress assays, we quantified the multi-trait heat tolerance of 569 individual colonies belonging to the *Acropora hyacinthus* species complex across 9.42° latitude of the Great Barrier Reef (GBR). We demonstrate considerable variation in the thermal thresholds of individual colonies both among (up to 7.3 °C) and within (up to 5.7 °C) reefs. Thermal history was the main driver of this variation, supporting the adaptation of populations to their local climatology (i.e., historical MMM) as well as acclimatization to recent thermal anomalies (i.e., DHW during collection). While Symbiodiniaceae community and the distribution of four host genomic clusters varied spatially, these factors had minimal contribution to variation in heat tolerance. Thus, unexplained variation in heat tolerance within putative species within the *A. hyacinthus* species complex may be controlled by functional genetic variation that was not measured in this study.

### Heat tolerance driven by thermal history

Sea surface temperature climatology was the strongest predictor of variation in the heat tolerance of colonies within the *A. hyacinthus* species complex across the GBR. This aligns with previous findings indicating that heat tolerance likely reflects local adaptation to temperature (Dixon et al., 2015; Howells et al., 2013; Kenkel & Matz, 2016; van Oppen et al., 2018). On northern and inshore reefs, *A. hyacinthus* individuals had higher thermal thresholds (F_v_/F_m_ and NDVI ED_50_s; **Fig. 2A-B**) but were living over 0.4°C closer to those thresholds than individuals on southern reefs. These populations may have higher absolute thermal thresholds but less capacity to adapt to future warming. While southern populations in this study exhibited lower absolute thermal thresholds, their greater distance from summer maximum temperatures indicates that they may be better positioned to persist through future warming, perhaps enhanced by gene flow of heat-adapted alleles from northern populations (Matz et al., 2018; Riginos et al., 2019). Similar latitudinal patterns in thermal thresholds have also been identified for *A. spathulata* in a parallel study (Denis et al., 2024) and in other non-acroporid species (Evensen et al., 2022; Howells et al., 2016; Marzonie et al., 2023; Woolsey et al., 2015), highlighting the potential for high latitude populations to serve as refugia as sea temperatures continue to rise.

In addition to climatology, recent accumulated thermal stress (i.e., DHW at time of collection) was an important predictor for two heat tolerance metrics (NDVI retention and F_v_/F_m_ ED_50_; **Fig. 6A,D**) . Colonies that were pre-exposed to mild to moderate recent heat stress (DHW > 0.5 and < 4) at the time of experimentation exhibited increased heat tolerance in our short-term assays. This supports that preconditioned colonies can display higher heat tolerance, likely via acclimatization (Brown et al., 2002; Thompson & van Woesik, 2009). This is an important consideration when using experimental heat stress to compare heat tolerance among colonies or sites as recent prior heat stress may lead to an overestimation of heat tolerance.

Further, more extreme heat stress would likely have opposite effects of mild heat stress and instead exacerbate damage to coral during subsequent experimental stress (Ainsworth et al., 2016; Denis et al., 2024). As heat tolerance varies with recent environmental conditions, acclimatization to these conditions may complicate understanding of heritable phenotypic variation available for natural or assisted evolution. Thus, future work should aim to reduce variation in heat stress preconditioning prior to assessing heat tolerance differences by performing experiments within the same season or including recent heat stress exposure in explanatory models.

### Bleaching variability among putative species within the Acropora hyacinthus complex

Analysis of genome-wide variation revealed four putative species in the *A. hyacinthus* complex on the GBR, with most samples representing the ‘neat’ morphotype, the primary target for this study (**Fig. 3**). This aligns with past work detecting four putative species within the *A. hyacinthus* complex on the GBR and six putative species across the Asia-Pacific (Ladner & Palumbi, 2012; Sheets et al., 2018). Our findings support the importance of a taxonomic revision to resolve morphological and genetic differences in this species complex.

Putative species displayed differences in their responses to a natural bleaching event, with Groups 2 and 3 exhibiting lower heat tolerance than Group 4 (**Fig. S4**). This suggests a role of host genomic identity in setting heat tolerance, although these differences may be confounded with low sample sizes of non-target putative species (n = 5, 7, and 31 for Groups 2 - 4). Further, it is possible that putative species within the *A. hyacinthus* complex hold distinct environmental preferences, as has been found in past work (Ladner & Palumbi, 2012; Rose et al., 2018; Suzuki et al., 2016). Thus, species effects on heat tolerance can be difficult to disentangle from other drivers (e.g., thermal history). Thus, in this study, our analysis of tolerance to experimental heat stress among genomic clusters was restricted to southern GBR colonies, which did not reveal differences among putative species, perhaps due to high trait variation within each genomic cluster. This conflicts with past work showing differences in natural bleaching responses and performance retention under experimental heat stress between two putative species within the *A. hyacinthus* complex living in the same backreef pool (Rose et al., 2018). Consequently, further work is needed to evaluate differences in heat tolerance among putative species within the *A. hyacinthus* complex while explicitly accounting for known genetic identities.

### High within-site variation in heat tolerance not well explained by predictors

For two of four heat tolerance metrics measured, we found variation among colonies within the *A. hyacinthus* complex was greater within reefs than among reefs (**Table 1; Table S3**), indicating that even small spatial scales may hold genetic variation in heat tolerance to allow adaptation to climate warming, even when accounting for host genomic cluster. This aligns with past work showing high trait variability among colonies within a single reef, including bleaching and mortality responses to experimental and natural heat stress (Cornwell et al., 2021; Cunning et al., 2016; Humanes et al., 2022). Here, thermal threshold variation in color paling (NDVI) was over 2.5 times more variable within sites compared to photochemistry (F_v_/F_m_). This may reflect the later measurement of this trait after the accumulation of more thermal damage (i.e., 17 versus 10 hours after the start of the experiment), or a more variable holobiont response in NDVI versus symbiont response in F_v_/F_m_. High phenotypic variability within small scales may reflect the high potential larval dispersal of *A. hyacinthus* as a broadcast spawner and may not reflect species with different life history strategies.

Predictors of heat tolerance were largely uninformative within reef sites, indicating that the high variability we measured among individual colonies is primarily determined by factor(s) other than depth, host genomic cluster identity, colony pigmentation, or Symbiodiniaceae community composition. Although we accounted for thermal and nutrient variables along depth stratifications, there likely remains additional unmeasured microhabitat features that could affect within-reef heat tolerance via acclimatization (Ainsworth et al., 2021). It is also likely that within-reef heat tolerance variation is controlled in part by host genetic variation given the high heritability of coral bleaching and survival under heat stress (reviewed in Bairos-Novak et al., 2021; Howells et al., 2022). Functional genomic variation within the coral host or Symbiodiniaceae may occur even across small scales (i.e., within a reef) and underpin heat tolerance variation that could not be accounted for in this study. Future work should aim to quantify this fine-scale genomic variation as it may be harnessed by natural or assisted evolution to support important restoration goals.

### Symbiodiniaceae communities were not major drivers of heat tolerance

Symbiodiniaceae communities in all *A. hyacinthus* clusters were composed of closely related and co-occurring variants of *Cladocopium* across the GBR. The predominant differentiation of Symbiodiniaceae communities by shelf position, followed by latitude, aligns with previous findings on the Symbiodiniaceae communities in acroporid corals on the GBR (Cooke et al., 2020; Cooper et al., 2011; Hoadley et al., 2019; Matias et al., 2022; Quigley et al., 2022; van Oppen et al., 2001). Inshore environments may exert unique selective pressure on Symbiodiniaceae communities due to higher temperatures, greater temperature or water variability, and/or increased turbidity (Cooke et al., 2020; Cooper et al., 2011; Matias et al., 2022; Tonk et al., 2013). We found minor differences in Symbiodiniaceae communities among different host genomic clusters, although this may be partially explained by unequal sampling of Groups 2, 3, and 4 in the southern GBR, where certain Symbiodiniaceae ITS2 variants were more abundant. Yet, this finding is consistent with past work showing that Symbiodiniaceae communities can vary between closely related putative host species (Johnston et al., 2022; Rose et al., 2021).

While Symbiodiniaceae communities varied spatially and by host cluster identity, they had minor effects on the heat tolerance of coral colonies in our study. This challenges previous findings showing that symbiont identity is a strong driver of heat tolerance in acroporid corals on the GBR; a pattern especially evident when coral colonies host multiple symbiont genera (Abrego et al., 2008; Jones et al., 2008). When limited to naturally-occurring variation in Symbiodiniaceae associations at the intrageneric level (e.g., within *Cladocopium*), heat tolerance conferred by the Symbiodiniaceae community may be less apparent. Yet, naturally and experimentally heat-adapted *Cladocopium* have been shown to enhance the heat tolerance of corals (Howells et al., 2012; Quigley et al., 2021), suggesting potential for heat tolerance variation among *Cladocopium* communities of *A. hyacinthus* across the GBR. Minor effects of Symbiodiniaceae on host heat tolerance detected in this study may be due to limitations of the ITS2 marker to detect species delineations and functional associations (Davies et al., 2023).

Additional markers such as psbA^ncr^ (LaJeunesse & Thornhill, 2011) may reveal a more complete picture of Symbiodiniaceae communities in these colonies, while genome-wide studies of adaptive variation (Ishida et al., 2023; Reich et al., 2021) may reveal insight into functional differences among closely related Symbiodiniaceae taxa and their effect on heat tolerance.

### Acute heat stress assay reveals colony-level variation

Our results follow recent applications of standardized acute heat stress experiments to quantify population thermal thresholds (Cunning et al., 2021; Evensen et al., 2022; Marzonie et al., 2023; Nielsen et al., 2022; Voolstra et al., 2021), extend this approach to colony-level applications (Cunning et al., 2021; Klepac et al., 2023), and is the first to quantify individual-level thresholds across environmental gradients in wild populations (but see Denis et al, 2024). This method allows rapid high-throughput phenotyping of corals and has been shown to measure relative differences in heat tolerance between sites and among individuals in a short term assay (18 hours) comparable to a longer-term assay (21 days; Klepac et al., 2023; Voolstra et al., 2020). To date, six other studies have been published using experimental aquaria to generate an ED_50_ metric based on F_v_/F_m_ measurements to compare thermal thresholds among populations (Evensen et al., 2022; Marzonie et al., 2023; Voolstra et al., 2021), among individuals in nursery-bred colonies (Cunning et al., 2021; Klepac et al., 2023), or among and within wild reef populations (Denis et al., 2024). Previous work has shown agreeability in heat tolerance rankings among nursery-bred colonies during two seasons in the same year (Cunning et al., 2021) and with bleaching resistance during a natural bleaching event (Morikawa & Palumbi, 2019), but further work is required to conclude that experimental heat tolerance responses are consistent across time and are indicative of survival during bleaching events.

Since no single trait can fully capture the coral heat tolerance phenotype, we measured both photochemical stress (the maximum quantum yield of photosystem II, F_v_/F_m_) and color paling (NDVI as a proxy for chlorophyll content) after acute heat stress. Recent work has shown that interpretations based on a single heat tolerance trait (i.e., bleaching vs photochemistry) after acute heat stress can lead to disparate conclusions, suggesting the importance of measuring multiple traits to yield a more comprehensive picture of heat tolerance (Klepac et al., 2023).

Future work should continue to assess multiple traits, especially to provide insight into both Symbiodiniaceae and host responses to heat stress. Standardized comparisons of heat tolerance utilizing multiple traits can allow for greater comparative frameworks across coral taxa and ocean regions.

## Conclusions

In this study, we find that heat tolerance within the *Acropora hyacinthus* species complex varies considerably across small and large spatial scales on the GBR. Depending on the heat tolerance trait measured, variation was sometimes greater within reef sites than among reef sites, suggesting that even small scales hold wide phenotypic variability within each putative species that can be harnessed by natural or artificial selection. Variation in heat tolerance rankings among traits suggests the importance in aligning tolerance metrics with research, restoration, or conservation goals. For example, absolute heat tolerance metrics may be of special interest to restoration initiatives aiming to enhance tolerance traits, while retained performance metrics may be more useful to understand how coral populations under climate warming will fare in their changing environments. Extensive variation in within-reef heat tolerance measured here indicates that heat tolerant alleles may be found across multiple reefs, supporting the notion that identifying heat tolerant broodstock for restoration is not as simple as sourcing from warm reefs (Howells et al., 2021; Quigley & van Oppen, 2022). Additional unexplained variation in heat tolerance may be due in part to functional genetic differences in the coral host or Symbiodiniaceae not measured here, which may uncover evolutionary potential in climate adaptation and standing variation available for restoration initiatives.

## Supporting information

Supplemental Methods and Results

## Acknowledgments

We thank Australian Institute of Marine Science (AIMS) technical staff and volunteers for field assistance, especially to R. Forester, C. Thompson, G. Cameron, S. Goyen, and M. Wooster. We also thank T. Bridge and A. Baird for assistance with species identification, C. Klass for image labelling, E. Hochberg and J. Kok for hyperspectral NDVI data extraction, M. Logan for statistical input, I. Byrne, C. Coppin, S. Howitt, Z. Meziere, and K. Prata for lab assistance, and C. Riginos for input on sample collection and sequencing. We acknowledge that Figures 1 and 3 were created with BioRender.com. We thank the Traditional Owners of the Great Barrier Reef for their free prior and informed consent to enter and undertake coral sampling in their Sea Country. We pay our respect to their Elders, past, present, and future.

## Author Contributions

EJH and LKB designed and funded the study. EJH, LKB, VMM, MSN, and HD conducted the field sampling and heat stress experiments. HD and MSN extracted the environmental variables and VMM extracted coordinate data. MSN extracted phenotypic data. PL and MSN conducted the ITS2 analysis. IP conducted the bioinformatic analysis and genomic cluster assignments.

MSN led the analysis of heat tolerance drivers. MSN wrote the manuscript with input from EJH. All authors contributed to and approved the final version of the manuscript.

## Data availability statement

The data and scripts that support the findings of this study are available at https://github.com/melissanaugle/Ahya_heat_tol_variation. Raw sequence data for species within the *Acropora hyacinthus* complex will be available under the NCBI Sequence Read Archive: PRJNA982441. Raw sequence data for Symbiodiniaceae will be made available in the NCBI Sequence Read Archive.

