## Supplemental Figures for "Environmental, host, and symbiont drivers of heat tolerance in a species complex of reef-building corals"

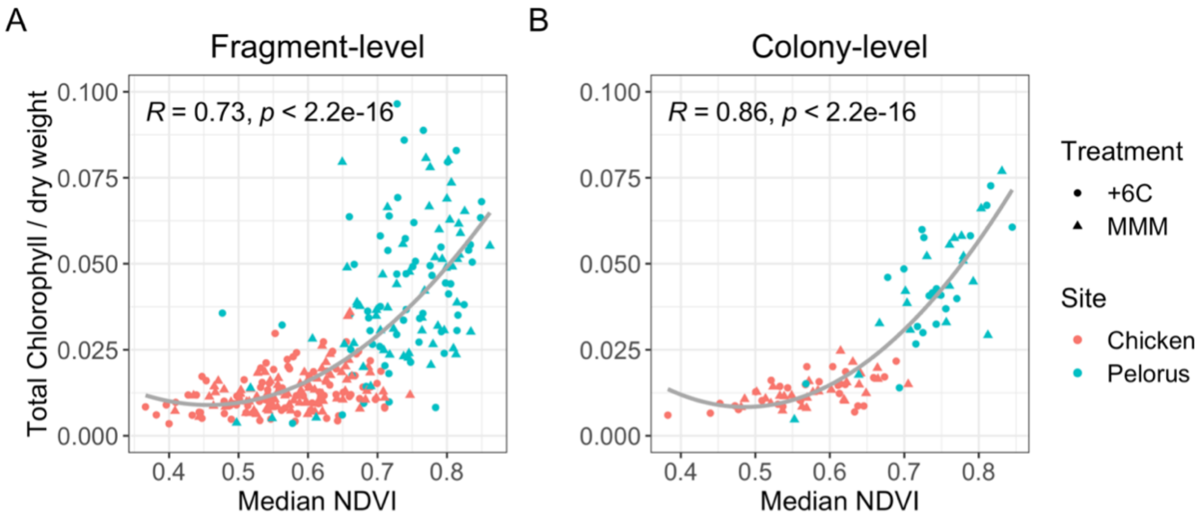


**Figure S1.** Relationship between hyperspectral normalized difference vegetation index (NDVI) and spectrophotometric total chlorophyll content in *Acropora hyacinthus* fragments (A) and colonies (B). *A. hyacinthus* fragments were from a subset of colonies from acute heat stress assays at Chicken (n=173) and Pelorus Island (n=120) following exposure to the local maximum monthly mean (MMM) and the MMM +6° C. R and p values represent the coefficients and statistical significance of Spearman correlations.

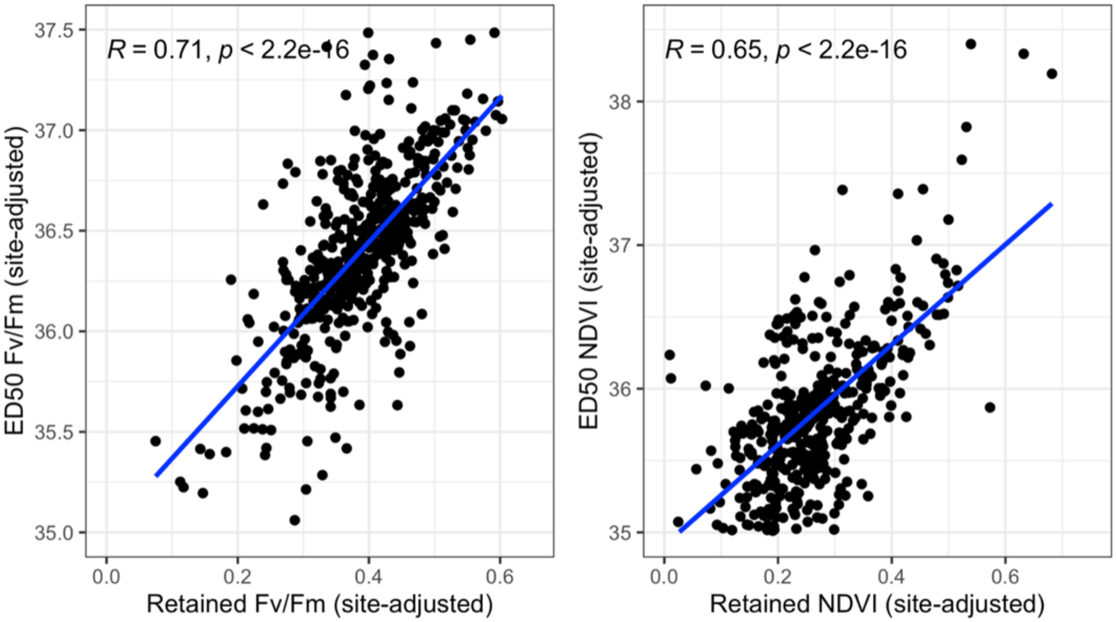


**Figure S2.** Relationships between site-adjusted metrics for $F_v/F_m$ (A) and normalised difference vegetation index (NDVI) (B) traits in *Acropora hyacinthus* colonies across the Great Barrier Reef. Site-adjusted absolute ($ED_{50}$) and retained performance (MMM +9°C/MMM) metrics are highly correlated when generated for the same trait, especially for $F_v/F_m$. *R* and *p* values indicate the strength and significance of Pearson correlations.

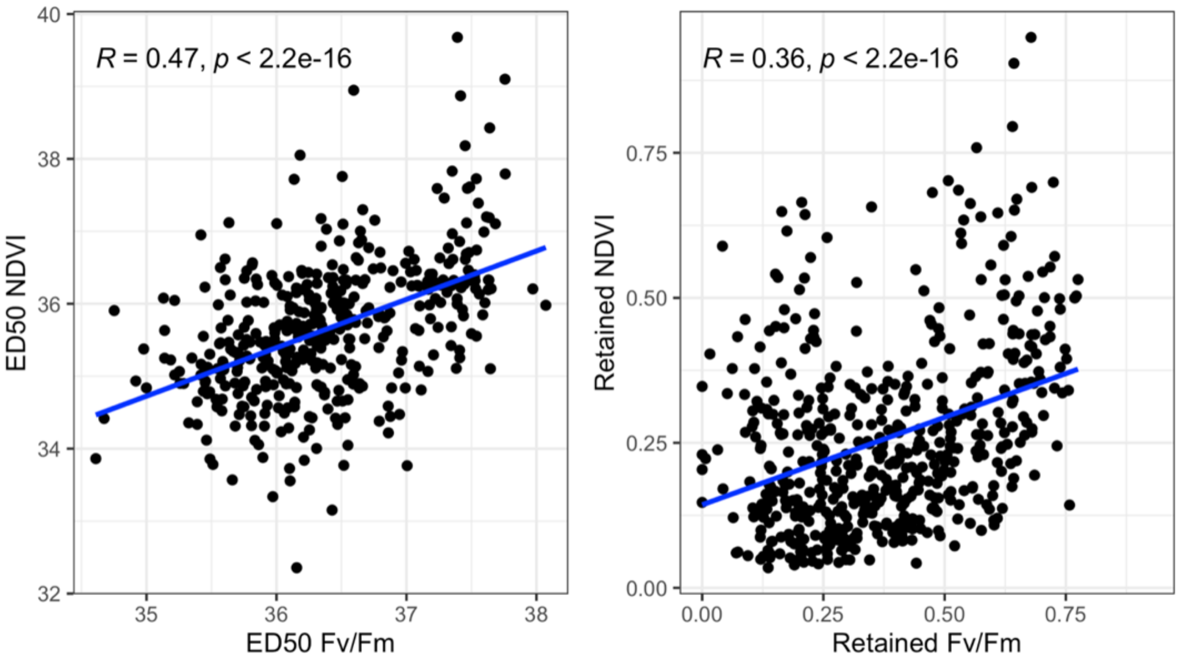


**Figure S3.** Relationships between thermal tolerance traits measured in *Acropora hyacinthus* colonies across the Great Barrier Reef. Moderate correlations between the maximum quantum yield of photosystem II ($F_v/F_m$) and the normalised difference vegetation index (NDVI) for colony values of absolute ($ED_{50}$; A) and retained (MMM +9°C/MMM; B) thermal tolerance metrics. *R* and *p* values indicate the strength and significance of Pearson correlations.

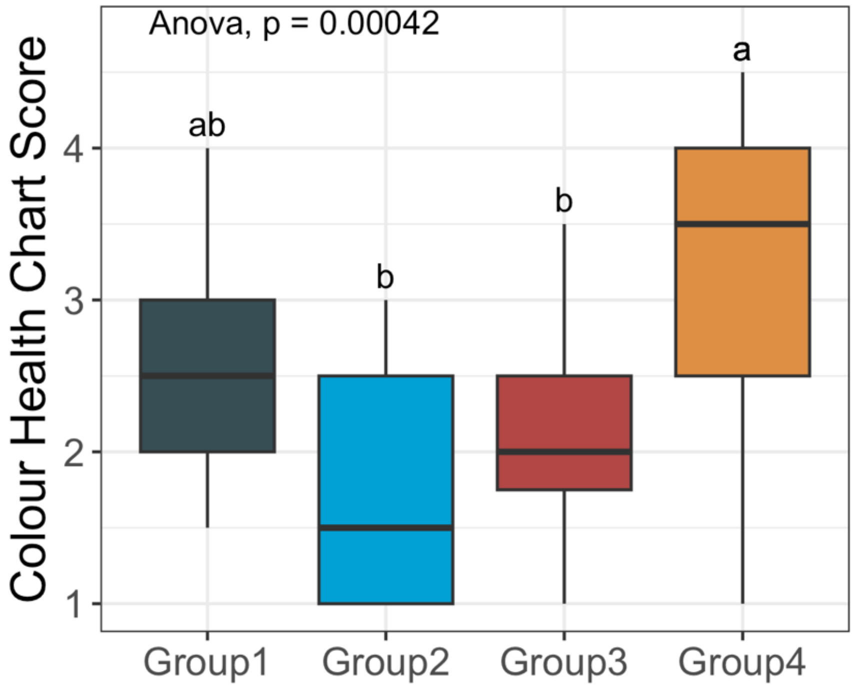


**Figure S4.** Variation in bleaching of *Acropora hyacinthus* (measured as CoralWatch Colour Health Chart Scores; Siebeck et al., 2006) by host genomic cluster in 60 colonies at North Direction on the GBR during a natural bleaching event in March 2021 (DHW ~ 3). Host genomic cluster was assigned based on genome-wide SNPs with Group 1 representing *A. hyacinthus* "neat" and Groups 2 - 4 representing other putative species in the *A. hyacinthus* complex. Sample sizes for Groups 1 – 4 were 17, 5, 7 and 31, respectively. Letters represent Tukey HSD post-hoc values from a one-way ANOVA.

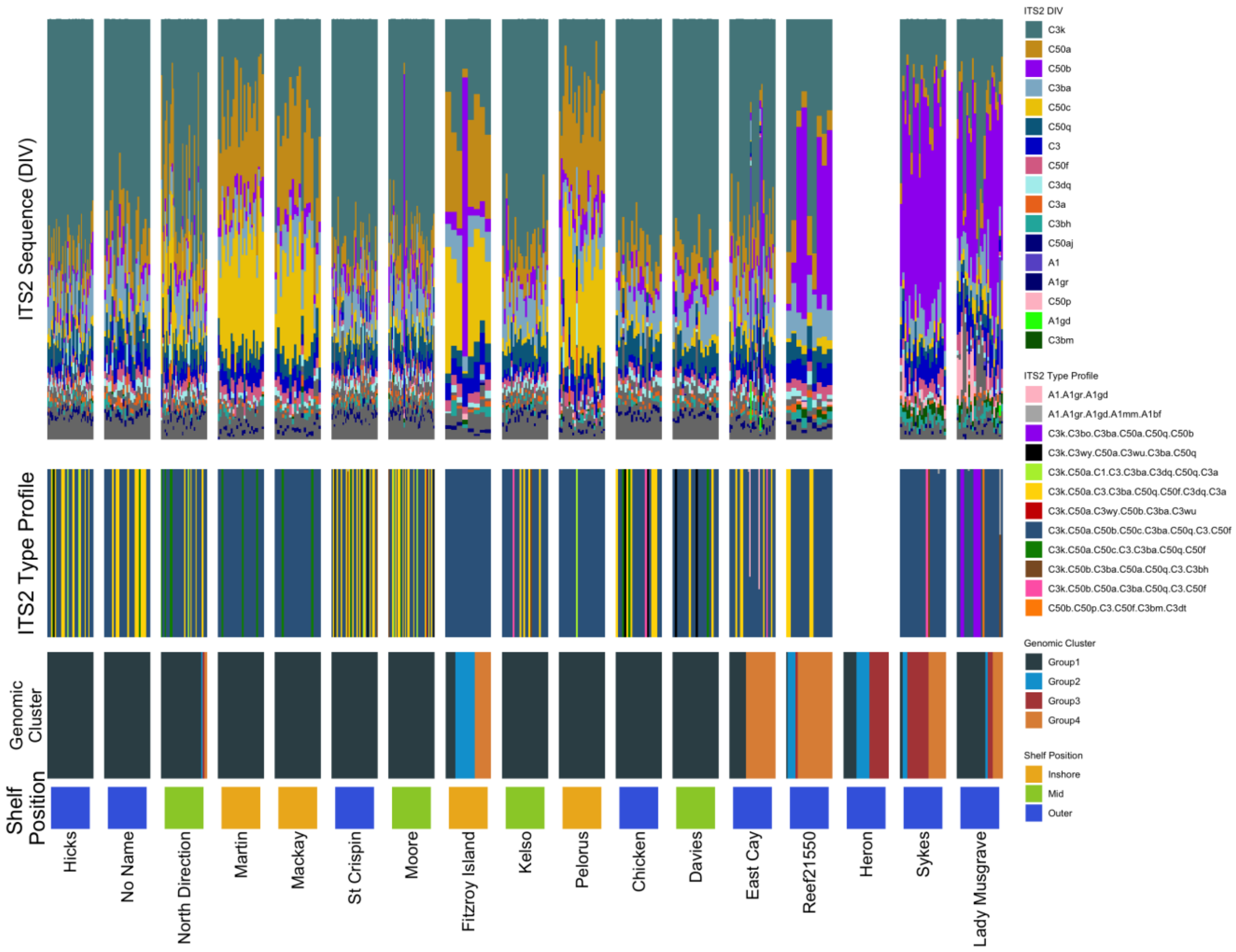


**Figure S5.** Variation in Symbiodiniaceae communities within *Acropora hyacinthus* colonies across the Great Barrier Reef. ITS2 sequences (top row), ITS2-type profiles (second row), and host genomic cluster (third row) are shown for individuals (vertical columns) within sites (arranged by latitude). Shelf position (bottom row) is shown to visualize ITS2 sequence differences at inshore sites (yellow, bottom row).

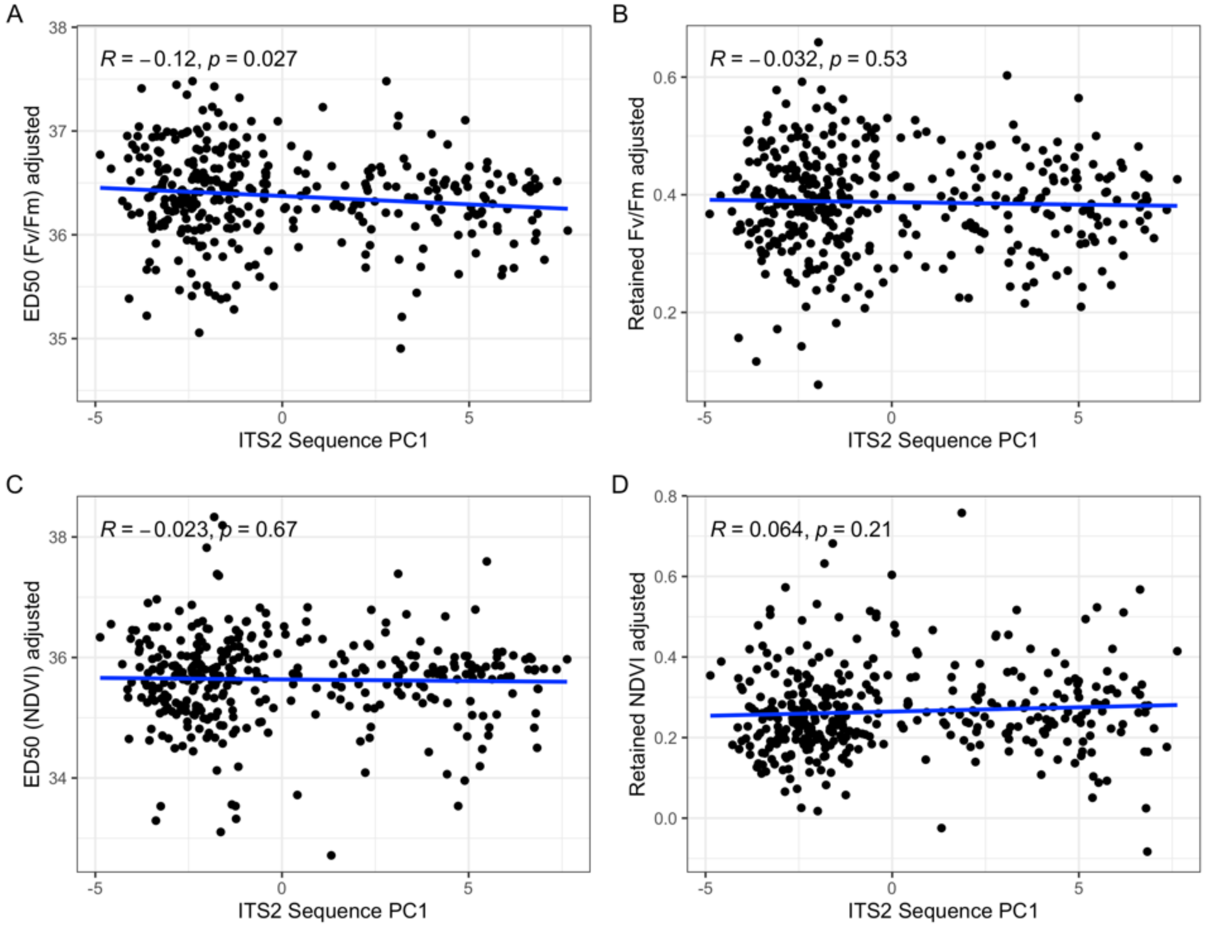


**Figure S6.** First PC value summarising 21.5% of the variation in Symbiodiniaceae ITS2 sequences and its association with heat tolerance traits in 400 colonies of *Acropora hyacinthus* from across the Great Barrier Reef. Heat tolerance trait metrics have been adjusted to remove variation due to site using residuals from site-specific means. Pearson correlation statistics are shown for each trait.

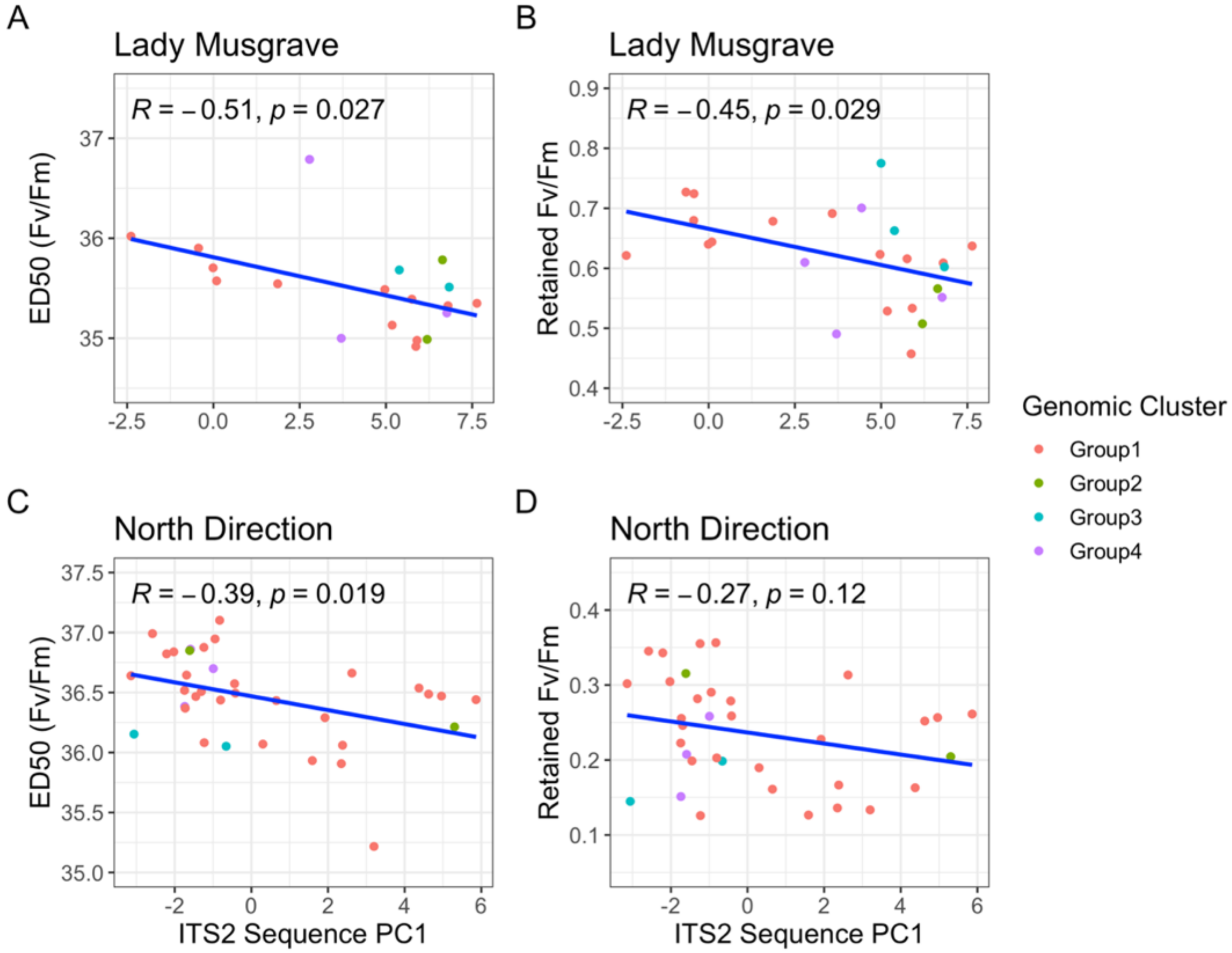


**Figure S7.** Relationships between Symbiodiniaceae ITS2 sequences and heat tolerance traits in *Acropora hyacinthus* colonies at Lady Musgrave (A, B) and North Direction (C, D) in the southern Great Barrier Reef. ITS2 PC1 exhibited moderate negative relationships with both $F_v/F_m$ derived metrics for Lady Musgrave (A, B) and for $F_v/F_m$ $ED_{50}$ for North Direction (C). Colonies are colored by genomic cluster to show the absence of a trend between genomic cluster and ITS2 sequence or heat tolerance.

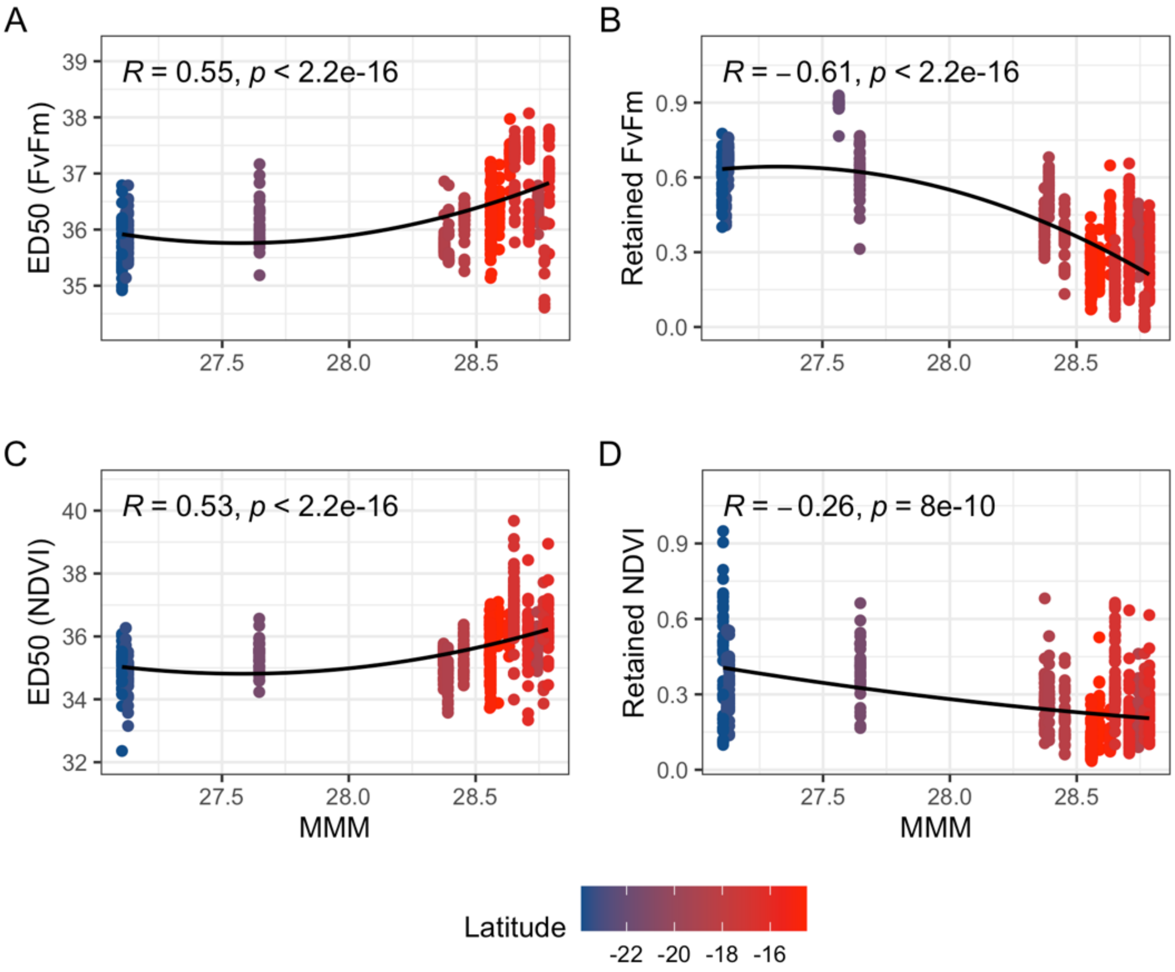


**Figure S8.** Relationships between the maximum monthly mean (MMM) and heat tolerance trait metrics for *Acropora hyacinthus* colonies on the Great Barrier Reef. All metrics demonstrate a significant relationship to MMM, which correlates with latitude. Relationships to MMM are positive for $ED_{50}$ thresholds of (A, C) and negative for retained performance metrics (B, D). Statistics represent Spearman correlations. ED50s with CIs < 5 were removed.

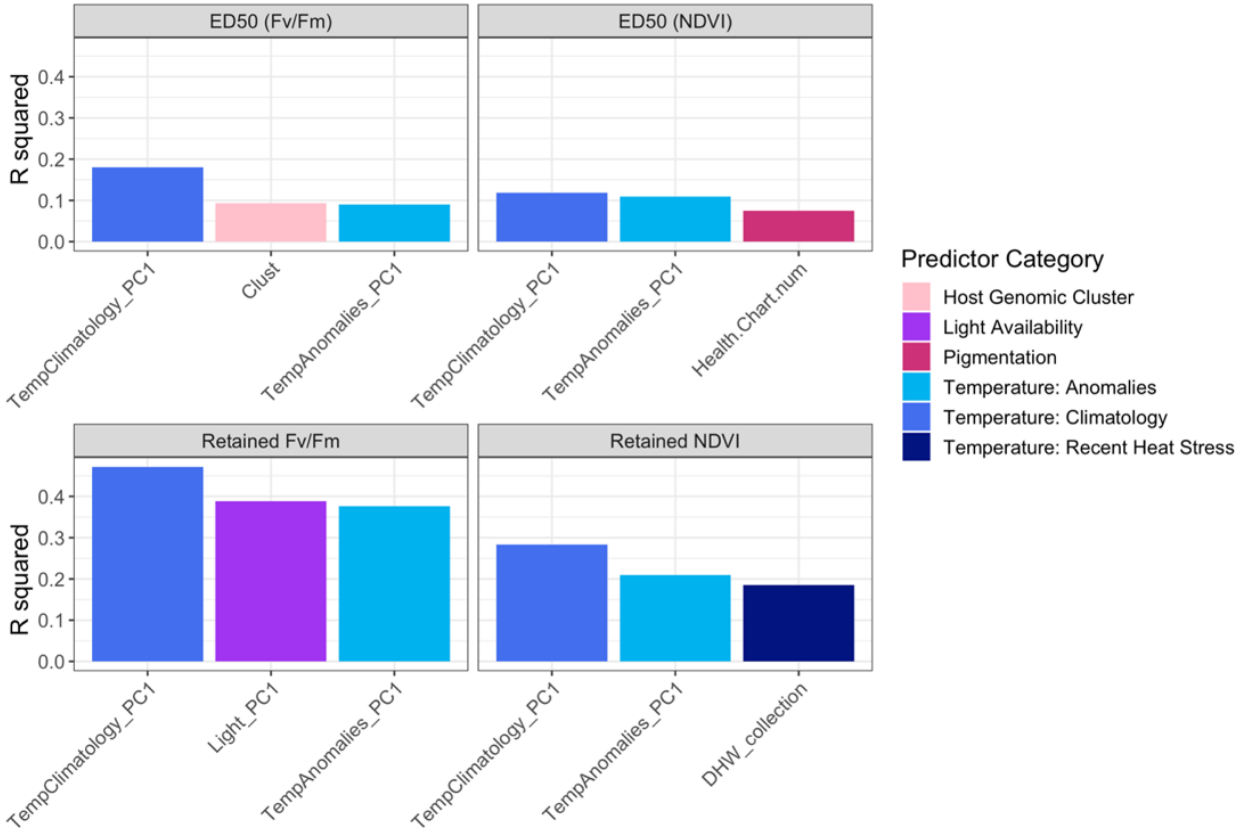


**Fig S9.** Relative influence of environmental predictors, as found through single-variable linear models, on the heat tolerance of *Acropora hyacinthus* on the Great Barrier Reef. $R^2$ values from the top 3 individual models for absolute thresholds ($ED_{50}$) and retained performance under heat stress for $F_v/F_m$ and NDVI, determined using AIC. Predictor categories represent PCA values summarizing variables falling within each functional category (**Table S7**).
