## Supplemental Methods and Results for "Environmental, host, and symbiont drivers of heat tolerance in a species complex of reef-building corals"

### Supplemental Information

### Methods

*ITS2 sequencing of Symbiodiniaceae*

Total DNA was extracted using a Qiagen Blood and Tissue Kit, including an overnight incubation followed by an RNase step with a 30-minute incubation. PCR amplification was performed using SYM_VAR_5.8S2 and SYM_VAR_REV primers (Hume et al. 2018) with Adapterama barcodes and Illumina adapters (Glenn et al., 2019). PCR reactions were performed with 6.25 µL Amplitaq Gold 360 Master Mix, 4.25 µL nuclease-free water, 0.5 µL forward primer (10 µM), 0.5 µL reverse primer (10 µM), and 1 µL DNA extract (10 ng/µL), totaling 12.5 µL. Thermocycler parameters were set as follows: 10 minutes at 95°C; 35 cycles at 95°C for 10 seconds, 56°C for 30 seconds, 72°C for 30 seconds; and a final cycle at 72°C for 5 minutes followed by storage at 4°C. PCR products were visualized on 1.5% agarose gels, with a clear single band of the expected length indicating successful amplification.

PCR products were bead-purified (PCR-DX) following the manufacturer's protocols. Products from independent plates were pooled in equimolar concentrations based on band intensity and placed in a limited-cycle PCR to facilitate the ligation of indexed iTru5 and iTru7 primers (Glenn et al., 2019). The second PCR step was performed in triplicate with 12.5 µL Amplitaq Gold 360 Master Mix (2x), 5 µL nuclease-free water, 1.25 µL forward primer (10 µM), 1.25 µL reverse primer (10 µM), and 5 µL pooled PCR 1 product, totaling 25 µL. Thermocycler parameters were set as follows: 10 minutes at 95°C; 12 cycles 95°C for 1 minute, 60°C for 30 seconds, and 72°C for 30 seconds; followed by 5 minutes at 72°C. Triplicates were pooled and bead-purified (PCR-DX) following the manufacturer's protocols. The final seven libraries were pooled in equimolar (per sample) proportions sequenced at the Australian Genome

Research Facility (Melbourne, Australia). Two replicate sequencing runs were performed across two lanes of an Illumina MiSeq 500 cycle with 250 bp paired-end reads. Barcoded samples were demultiplexed using `MrDemuxy` version 1.2.0 in Python 2.7 (https://github.com/lefeverde/Mr_Demuxy). Read number and sequence quality were assessed using `FastQC` (http://www.bioinformatics.babraham.ac.uk/projects/fastqc/).

#### *Ranking Heat Tolerance Predictors using Linear Modelling*

Predictive ability of environmental, Symbiodiniaceae, and host genetic identity variables on heat tolerance were assessed using additional statistical methods to support findings of the Boosted Regression Trees (BRTs) stated in the main text. Linear models and stepwise multiple regression were used to investigate effects of predictors on heat tolerance traits. Since numerous correlated predictors were used in the BRTs, variables within a similar category were summarized using PCA to reduce the predictor set to 10 (**Table S7**). Of these 10 predictors, only combinations of predictors with $R^2 < 0.8$ were used in the same model to reduce collinearity. Stepwise linear model selection using the stats R package was used to retain only variables that lowered model AIC score. To understand individual contributions of predictor categories on heat tolerance traits, individual linear models were built and adjusted $R^2$ values were compared.

### Results

#### *Linear Models Support Predictive Ability of Thermal History on Heat Tolerance*

The importance of thermal history variables as predictors of thermal tolerance, as found in the BRTs, was confirmed using linear models for each predictor category (**Fig. S9**). Temperature climatology PC1, characterised by MMM, overall mean temperature, and lower monthly mean temperature (**Table S6**), was identified as the strongest predictor for all heat tolerance metrics ($R^2$= 0.12 to 0.53) as in the BRT framework. Similarly, temperature anomaly PC1 ($R^2$= 0.09 to 0.50) was among the top fitting models across all metrics. Genomic cluster and light variables also held some predictive power for $F_v/F_m$ metrics (**Fig. S9**), which may reflect their strong correlation with temperature climatology or the influence of light on photochemical traits. Recent heat stress, measured as DHW at the time of collection, was informative in explained retained NDVI (**Fig. S9**; $R_2$ = 0.19), agreeing with the BRT model. CoralWatch Health Chart Score measured during collection was minimally informative in explaining NDVI $ED_{50}$ ($R_2$ = 0.08) and did not correlate with latitude or depth. In stepwise multiple regression, temperature climatology and host genomic cluster were retained in all models (**Table S6**). CoralWatch Health Score was retained in 3 of 4 final models (**Table S6**).

Variance explained for each trait aligned among BRTs, linear models using predictor categories, and full linear models, with $F_v/F_m$ better predicted than NDVI and retained performance better predicted than $ED_{50}$ **(Fig. 6; Table S6; Fig S9).** Full linear model adjusted $R^2$ values were similar to BRT deviance explained for retained NDVI (adj. $R^2$ = 0.46, deviance explained = 0.48), but held less predictive ability for NDVI $ED_{50}$ (adj. $R^2$ = 0.20, deviance explained = 0.37), $F_v/F_m$ $ED_{50}$ (adj. $R^2$ = 0.24, deviance explained = 0.67) and retained $F_v/F_m$ (adj. $R^2$ = 0.58, deviance explained = 0.81; **Table S7**; **Fig. 6**).
