## Supplemental Tables for "Environmental, host, and symbiont drivers of heat tolerance in a species complex of reef-building corals"

**Table S1.** *Acropora hyacinthus* colonies sampled from the Great Barrier Reef for evaluating phenotypic variation in thermal tolerance using acute heat stress assays. 60 additional colonies were sampled at North Direction during a bleaching event in 2021 but were not evaluated in the acute heat stress assay.

| Reef Site | Latitude | Longitude | Date(s) Sampled | n Colonies |
|---|---|---|---|---|
| Reef 21550 | -21.9455 | 152.3135 | 19 & 20 Jan. 2021 | 29 |
| East Cay | -21.4960 | 152.5560 | 21 & 22 Jan. 2021 | 33 |
| Lady Musgrave | -23.8929 | 152.4064 | 25 Jan. 2021; 24 March 2022 | 42 |
| Sykes | -23.4459 | 152.0244 | 26 Jan. 2021 | 32 |
| Hicks | -14.4735 | 145.5046 | 19 & 20 March 2021 | 43 |
| North Direction | -14.7434 | 145.5133 | 22 & 23 March 2021 | 53 |
| No Name | -14.6508 | 145.6383 | 25 March 2021 | 34 |
| Martin | -14.7801 | 145.3681 | 27 March 2021 | 30 |
| St Crispin | -16.0724 | 145.8449 | 29 March 2021; 15 April 2021 | 44 |
| Kelso | -18.4240 | 146.9849 | 11 April 2021 | 33 |
| Pelorus East | -18.5587 | 146.5023 | 12 April 2021 | 30 |
| Mackay | -16.0384 | 145.6514 | 16 April 2021; 4 March 2022 | 43 |
| Chicken | -18.6706 | 147.7064 | 20 April 2021 | 29 |
| Davies | -18.8270 | 147.6268 | 21 April 2021 | 30 |
| Fitzroy Island | -16.9406 | 145.9948 | 6 May 2021 | 14 |
| Moore | -16.8457 | 146.2316 | 7 May 2021 | 50 |
| Heron Island | -23.4416 | 151.9130 | 26 March 2022 | 14 |
| **Total** | | | **26 experimental assays** | **583** |

**Table S2.** Predictors used in the Boosted Regression Tree analysis of phenotypic variation in the heat tolerance of *Acropora hyacinthus* across the Great Barrier Reef. Temperature variables of climatology, anomalies, and recent heat stress were calculated using CoralWatch data since they rely on DHW which corresponds to the historical MMM which is calculated using the CoralWatch dataset. Temperature metrics of variability were calculated using eReefs, which provides higher resolution recent temperature data.

| Identifier | Variable Type | Description |
|---|---|---|
| Latitude | Location | Latitude |
| Shelf.pos | Location | Shelf Position (i.e., inshore, mid, offshore) |
| MMM | Temp: Climatology | MMM (adjusted to historical baseline) |
| TSA_DHW_mean | Temp: Anomalies | Mean DHW |
| TSA_DHW_stdev | Temp: Anomalies | DHW SD |
| DHW_freq_sup4 | Temp: Anomalies | Mean frequency of DHW > 4 |
| DHW_freq_sup8 | Temp: Anomalies | Mean frequency of DHW > 8 |
| DHW_max | Temp: Anomalies | Maximum DHW |
| SSTA_freq_stdev | Temp: Anomalies | SD of temp. anomalies |
| SSTA_freq_mean | Temp: Anomalies | Mean temp. anomalies |
| DTR | Temp: Variability | Daily thermal range mean |
| DTR_ss | Temp: Variability | Daily thermal range (spring-summer mean) |
| AR | Temp: Variability | Mean annual range |
| ROTC_ss | Temp: Variability | Mean rate of temp. change (spring-summer) |
| DHW_collection | Temp: Recent heat stress | DHW at the time of collection |
| Depth | Depth | Colony depth (adjusted to LAT using BOM tide data) |
| Health.Chart.Score | Pigmentation | CoralWatch Colour Health Chart Score at collection |
| CF_OM | Light Availability | Cloud frac. Overall mean |
| Secchi_OM | Light Availability | Secchi from 488nm mean |
| TOTAL_NITROGEN_OM | Nutrient Availability | Total nitrogen mean |
| NO3_OM | Nutrient Availability | Nitrate mean |
| PIP_OM | Nutrient Availability | Particulate inorganic phosphorus mean |
| ITS2variant_PC1 | Symbiont Community | First PC value summarizing all ITS2 sequences |
| ITS2variant_PC2 | Symbiont Community | Second PC value summarizing all ITS2 sequences |
| Dominant_type | Symbiont Community | Dominant ITS2 type profile |
| Clust | Host Genomic Cluster | Assignment to one of four host genetic clusters |

**Table S3.** Linear mixed effects models (LMMs), accounting for genomic cluster identity, showing the among-site variation and residual (within-site) variation of heat tolerance trait metrics in colonies within the *Acropora hyacinthus* complex across sites on the Great Barrier Reef. Higher within-site variance was exhibited by the normalised difference vegetation index (NDVI) absolute metric ($ED_{50}$ threshold; A) and retained performance metric (B); while greater among site variance was exhibited by the maximum quantum yield of photosystem II ($F_v/F_m$) absolute metric ($ED_{50}$ threshold; C) and retained performance metric (D).

| | | **Variance** | **Standard Deviation** |
|---|---|---|---|
| A. $F_v/F_m$ $ED_{50}$ | Site | 0.3017 | 0.5493 |
| | Residual | 0.1883 | 0.4340 |
| | Genomic Cluster | 0.0054 | 0.0735 |
| B. Retained $F_v/F_m$ | Site | 0.0434 | 0.2084 |
| | Residual | 0.0072 | 0.0851 |
| | Genomic Cluster | 0.0001 | 0.0104 |
| C. NDVI $ED_{50}$ | Site | 0.3494 | 0.5911 |
| | Residual | 0.5189 | 0.7204 |
| | Genomic Cluster | 0.0590 | 0.2429 |
| D. Retained NDVI | Site | 0.0105 | 0.1025 |
| | Residual | 0.0136 | 0.1168 |
| | Genomic Cluster | 0.000 | 0.000 |

**Table S4.** PERMANOVA table showing the effect of latitude, shelf position, host genomic cluster, and their interactions on the composition of Symbiodiniaceae ITS2 sequence variants in 398 *Acropora hyacinthus* colonies from across the Great Barrier Reef. ITS2 sequence diversity PC values were influenced by all factors, but most strongly by shelf position, followed by latitude (partial $\omega^2$ = 0.4100, 0.2312).

| | df | $R^2$ | F | Partial $\omega^2$ | *p-value* |
|---|---|---|---|---|---|
| Shelf_position | 2 | 0.3044 | 139.271 | 0.4100 | 0.001 |
| Latitude | 1 | 0.1319 | 120.670 | 0.2312 | 0.001 |
| Host_cluster | 3 | 0.0464 | 14.148 | 0.0902 | 0.001 |
| Shelf_position:Latitude | 2 | 0.0366 | 16.746 | 0.0733 | 0.001 |
| Shelf_position:Host_cluster | 5 | 0.0176 | 3.221 | 0.0271 | 0.001 |
| Latitude:Host_cluster | 3 | 0.0468 | 14.277 | 0.0910 | 0.001 |
| Residual | 381 | 0.4164 | | | |
| Total | 397 | 1.0000 | | | |

**Table S5.** PERMANOVA table showing the effect of latitude, shelf position, host genomic cluster, and their interactions on Symbiodiniaceae ITS2 type profiles in 403 *Acropora hyacinthus* colonies from across the Great Barrier Reef. ITS2 type profiles were most influenced by shelf position and latitude (p = 0.001).

| | df | $R^2$ | F | Partial $\omega^2$ | *p-value* |
|---|---|---|---|---|---|
| Shelf Position | 2 | 0.0390 | 8.3049 | 0.0350 | 0.001 |
| Latitude | 1 | 0.0272 | 11.598 | 0.0256 | 0.001 |
| Host_cluster | 3 | 0.0132 | 1.8717 | 0.0064 | 0.077 |
| Shelf_position:Latitude | 2 | 0.0052 | 1.1089 | 0.0005 | 0.339 |
| Shelf_position:Host_cluster | 5 | 0.0069 | 0.5890 | -0.0051 | 0.848 |
| Latitude:Host_cluster | 3 | 0.0026 | 0.3707 | -0.0047 | 0.868 |
| Residual | 386 | 0.9060 | | | |
| Total | 402 | 1.0000 | | | |

**Table S6.** Linear models of drivers of heat tolerance traits in *Acropora hyacinthus* on the Great Barrier Reef as determined through stepwise model selection using AIC. SE represents standard error.

| | | **Estimate** | **SE** | **tvalue** | **pvalue** |
|---|---|---|---|---|---|
| **Fv/Fm ED50 (adj $R^2$ = 0.234)** | Intercept | 36.10 | 0.12 | 297.11 | 0.000 |
| | Temp. climatology PC1 | -0.19 | 0.03 | -5.53 | 0.000 |
| | Depth | -0.09 | 0.04 | -2.39 | 0.018 |
| | Genetic cluster group 2 | -0.61 | 0.21 | -2.86 | 0.005 |
| | Genetic cluster group 3 | 0.20 | 0.22 | 0.90 | 0.368 |
| | Genetic cluster group 4 | 0.01 | 0.19 | 0.07 | 0.943 |
| | ITS2 variant PC1 | 0.05 | 0.02 | 3.18 | 0.002 |
| | Coral health chart score | 0.09 | 0.04 | 2.54 | 0.018 |
| | Nutrients_PC1 | 0.04 | 0.02 | 2.46 | 0.015 |
| **Retained Fv/Fm (adj $R^2$ = 0.580)** | Intercept | 0.46 | 0.03 | 18.77 | 0.000 |
| | Temp. climatology PC1 | 0.07 | 0.01 | 8.78 | 0.000 |
| | Depth | -0.02 | 0.01 | -2.72 | 0.007 |
| | Genetic cluster group 2 | -0.14 | 0.04 | -3.86 | 0.000 |
| | Genetic cluster group 3 | -0.04 | 0.04 | -0.93 | 0.355 |
| | Genetic cluster group 4 | -0.11 | 0.03 | -3.28 | 0.001 |
| | Coral health chart score | -0.01 | 0.01 | -2.27 | 0.024 |
| | Temp. anomalies PC1 | 0.02 | 0.01 | -2.61 | 0.010 |
| | DHW during collection | 0.01 | 0.01 | 1.50 | 0.136 |
| **NDVI ED50 (adj $R^2$ = 0.200)** | Intercept | 34.80 | 0.17 | 204.99 | 0.000 |
| | Temp. climatology PC1 | -0.021 | 0.05 | -4.55 | 0.000 |
| | Coral health chart score | 0.024 | 0.05 | 5.24 | 0.000 |
| | Genetic cluster group 2 | -0.97 | 0.32 | -3.09 | 0.002 |
| | Genetic cluster group 3 | 0.00 | 0.32 | 0.01 | 0.989 |
| | Genetic cluster group 4 | 0.16 | 0.27 | 0.59 | 0.559 |
| **Retained NDVI (adj $R^2$ = 0.465)** | Intercept | 0.22 | 0.02 | 9.01 | 0.000 |
| | Temp. climatology PC1 | 0.03 | 0.01 | -4.84 | 0.000 |
| | DHW during Collection | -0.05 | 0.01 | -7.66 | 0.000 |
| | Temp. variability PC1 | -0.04 | 0.01 | -5.45 | 0.000 |
| | Coral health chart score | 0.03 | 0.01 | 5.01 | 0.000 |
| | Depth | -0.03 | 0.01 | -3.07 | 0.002 |

**Table S7.** Predictors used to generate PC values used in multiple regression and linear models of phenotypic variation in the heat tolerance of *Acropora hyacinthus* across the Great Barrier Reef. PIP represents particulate inorganic phosphorus. KD represents vertical attenuation.

| PC category | Identifier | Description |
|---|---|---|
| Anomalies | TSA_DHW_mean | Mean DHW |
| Anomalies | TSA_DHW_stdev | DHW SD |
| Anomalies | DHW_freq_sup4 | Mean frequency of DHW > 4 |
| Anomalies | DHW_freq_sup8 | Mean frequency of DHW > 8 |
| Anomalies | DHW_max | Maximum DHW |
| Anomalies | SSTA_freq_stdev | SD of temp. anomalies |
| Anomalies | SSTA_freq_mean | Mean temp. anomalies |
| Climatology | MMM | MMM (adjusted to historical baseline) |
| Climatology | OM | Overall mean |
| Climatology | LMM | Lower monthly mean (adjusted to historical baseline) |
| Temp. Variability | DTR_ss | Daily thermal range mean |
| Temp. Variability | DTR | Daily thermal range (spring-summer mean) |
| Temp. Variability | ROTC_ss | Mean rate of temp. change (spring-summer) |
| Temp. Variability | AR | Mean annual range |
| DHW_collection | DHW_collection | DHW at the time of collection |
| Nutrients | Chl_a_HMsd | SD of high monthly mean of chlorophyll a |
| Nutrients | Chl_a_HM | High monthly mean of chlorophyll a |
| Nutrients | Chl_a_OMsd | SD of chlorophyll a |
| Nutrients | Chl_a_OM | Mean chlorophyll a |
| Nutrients | TOTAL_NITROGEN_HMsd | SD of high monthly mean of total nitrogen |
| Nutrients | TOTAL_NITROGEN_HM | High monthly mean of total nitrogen |
| Nutrients | TOTAL_NITROGEN_OMsd | SD of total nitrogen |
| Nutrients | TOTAL_NITROGEN_OM | Mean total nitrogen |
| Nutrients | PIP_HMsd | SD of high monthly mean of PIP |
| Nutrients | PIP_HM | High monthly mean of PIP |
| Nutrients | PIP_OMsd | SD of PIP |
| Nutrients | PIP_OM | Mean PIP |
| Nutrients | NO3_HMsd | SD of high monthly mean of nitrate |
| Nutrients | NO3_HM | High monthly mean of nitrate |
| Nutrients | NO3_OMsd | SD of nitrate |
| Nutrients | NO3_OM | Mean nitrate |
| Light | Kd_490_HM | SD of high monthly mean of KD at 490 nm |
| Light | Kd_490_HMsd | High monthly mean of KD at 490 nm |
| Light | Kd_490_OMsd | SD of KD at 490 nm |
| Light | Kd_490_OM | Mean KD at 490 nm |
| Light | CF_OM | Mean cloud fraction |
| Light | CF_ss | Mean cloud fraction during spring-summer |
| Light | Secchi_HMsd | SD of high monthly mean of secchi depth |
| Light | Secchi_HM | High monthly mean of secchi depth |
| Light | Secchi_OMsd | SD of secchi depth |

| | | |
|---|---|---|
| Light | Secchi_OM | Mean secchi depth |
| ITS2.variant | ITS2 variants | Symbiodiniaceae ITS2 sequence diversity |
| Clust | Clust | Host genomic cluster |
| Depth.corrected | Depth | Colony depth |
| Health.chart.num | Health.chart | CoralWatch Health Chart Score |
